# Sustained quenching not always means photoinhibition

**DOI:** 10.1101/2025.09.03.673506

**Authors:** Maximiliano Cainzos, Maria Dolores Pissolato, Chen Hu, Nazeer Fataftah, Sanchali Nanda, Stefan Jansson

## Abstract

Photosynthetic light harvesting complexes (LHC) are involved in light absorption and energy dissipation. By modulating the photosystems absorption cross section, they affect their photosynthetic activity and non-photochemical quenching (NPQ) capacity. These processes have been widely studied by spectrally integrated chlorophyll fluorescence methods, which mask their associated spectral information. We explored in aspen and Arabidopsis npq mutants how the absence of these components affects the development of NPQ spectra under two contrasting conditions: in the absence and presence of photoinhibition. We proposed a new parameter to estimate the development of new emitting species (NESD) during time-spectrally resolved NPQ inductions and a pipeline to disentangle PSII energy partitioning heterogeneity. We demonstrate that LHCB, PsbS and zeaxanthin is required for NESD. By combining gas exchange with spectrally resolved kinetics, we show that under photoinhibitory conditions, however, NES develops in the absence of PsbS and zeaxanthin, and the resulting sustained quenching occurring independently of photoinhibition. Furthermore, we found that in the absence of LHCB and Curvature Thylakoid 1 a significant increase in photoinhibition was observed. This suggest that in the long term effective photoprotection requires the presence of LHCB and thylakoid plasticity, while PsbS and zeaxanthin play a major role in catalyzing LHCII-dependent quenching.

## Introduction

Light harvesting is a fundamental process in green photosynthetic organisms, taking place in photosystem II (PSII) and photosystem I (PSI) in the thylakoid membrane. To harvest and efficiently convert solar energy into chemical energy, photosystems have a matrix of pigments excited by light and transferring excitation energy to reaction centers (RC) where charge separation takes place (for review, see Croce and Van Amerongen 2020). During evolution, Chlorophytes have expanded their photosystems by evolving light harvesting complexes, increasing the absorption cross section of PSII and PSI by binding efficiently connected pigments to their RC (LHC; Bag 2021). *In planta*, the structure of PSII and PSI core is composed of chlorophylls a and carotenoids while their LHC contain chlorophylls a, b and carotenoids. Due to its nature, light is constantly shifting its spectrum and intensity, making the LHC important for the fine-tuning of energy transfer to their RC. In addition to their harvesting function, LHCs participate in photoprotection, in which LHCII associated with PSII makes an important contribution (Ruban and Saccon 2022).

Non-photochemical quenching (NPQ) can be described as the general photoprotection mechanism that reduces the rate of excitation energy transfer (EET) to the PSII RC, thereby preventing RC overreduction and ROS production (Ruban, 2016). Several NPQ (sub)processes have been described/characterized, from the fast PsbS-dependent response (qE) to slow “sustained quenching”. Since “sustained” refers only to the kinetics, it remains unclear how many distinct molecular mechanisms contribute to this type of quenching. Mechanism(s) dependent on Zea or lipocalin have been denoted as qZ and qH, respectively (for review see Malnoë, 2018) but direct energy transfer from PSII to PSI has also in some systems been shown to provide sustained quenching (see e.g. Bag et al. 2020). In the absence of qZ it has been assumed that the slow arise of sustained quenching is solely the product of photoinhibition, which is referred as qI (Nilkens et al., 2010; Ramakers et al., 2024). Under that situation, it is expected that increasing sustained quenching should be followed by a strong decline in the photochemistry.

Under NPQ conditions, a long-wavelength fluorescence shoulder (peaking around 720– 740 nm, Farooq et al. 2018) is observed when PSII RC are closed, hereafter referred as the new emitting species (NES). Crucially, this spectral signature has also been linked to photoprotection, whereas its absence is associated with the lack of qE (Holzwarth et al., 2009). Since its discovery in the early 1990s, NES has been attributed to different processes. Originally, it was postulated that qE quenching and LHCII aggregation is the source of the NES and is dependent of PsbS, where Zea enhances this spectral property (Horton et al. 1991; Holzwarth et al. 2009; Johnson and Ruban, 2009). Although this signature has been associated to LHCII quenching, also it can be largely affected by PSII-PSI energy transfer in gymnosperms, where the NPQ spectra has a strong PSI enhanced emission in combination with the contribution of LHCII quenching (Bag et al., 2020). However, recent reports have challenged this interpretation and proposed that the NES originates from PSII closed RC under actinic light, where dynamic feedback at PSII RC contributes to photoprotection (Farooq et al. 2018).

One way to address these uncertainties would be to study the NPQ spectral properties of PSII RC closed in the absence and presence of PSII antenna and truncated light harvesting antenna (TLA) mutants are therefore an excellent resource for this purpose. Since LHCIIs are not essential for the plant and do not affect the functionality of the RC, several strategies have been developed to study how the photosynthetic machinery behaves in their absence (Cutolo et al. 2023). One is to reduce the physical PSII antenna size by the manipulation of the chlorophyll b content and chlorophyll a oxygenase mutants have therefore been used to reduce the LHCII content in plants, as chl b is essential for the stability of several of the LHCs. However, until now the spectral properties of fluorescence inductions and NPQ in TLA plants have not been studied.

In recent years, a large effort has been made to enhance photosynthetic productivity in crops to meet the standard for worldwide security (Leister, 2023). However, photosynthesis improvement has been largely unexplored in trees which offers the opportunity of boost plant based second-generation biofuels productivity, like in fast-growing trees such as aspen (Carriquiri et al., 2011). To study some of the key components involved in photosynthetic regulation in aspen, we developed a collection of chlorophyll a oxygenase (*chlorina*) mutants, which was combined with aspen NPQ mutants where PsbS (*psbs*) or Violaxanthin De-Epoxidase (*vde*) is missing (Nanda et al., 2025). From them, we disentangle some of the unknown spectral properties associated to NPQ. We were able to assess the contributions of PsbS, Zea, and LHCB to the heterogeneous development of the NPQ spectrum during the initial phase of NPQ. Additionally, for the first time, we investigated the impact of NPQ on the wavelength-dependent energy partitioning of PSII, providing new insights into this process. We found different spectral properties for PSII energy partitioning in classical photosynthetic mutants, suggesting a complex interplay between leaf light gradient and NPQ. Lastly, by performing photoinhibitory treatments monitored in real-time by gas exchange and chlorophyll a fluorescence, we disentangle some of the components required for photoprotection and their relationship with sustained quenching, photoinhibition and the development of NES *in folio*. Due to the unexpected outcome, we conducted a comparative analysis using well-known Arabidopsis mutants to determine the extent to which the universality of these findings applies, an aspect that has gained increasing relevance in recent years (Leister, 2023).

## Material and Methods

### Plant material

Aspen and *Arabidopsis* were grown at 22°C/16°C, 16hs/8hs day/night cycle for up to 12 weeks (Supplementary Fig. S1), light intensity was 150-180 μmol photons m^−2^ s^−1^. Aspen npq mutants for PsbS (*psbs*) and Violaxanthin-De-Epoxidase (*vde*) were included (Nanda et al., 2025) in combination with Chlorophyll a oxygenase (CAO) mutants. Arabidopsis mutants affecting NPQ key players were used, including *npq1* (lacking VDE), *npq4* (lacking PsbS), L17 (overexpressing PsbS) and *curt1* (CURVATURE THYLAKOID 1 proteins; Havaux and Nigoyi, 1999; Li et al., 2000; Li et al., 2002; Armbruster et al., 2013).

### Design and cloning of CRISPR-single guide RNAs (sgRNAs) constructs

Two genes potentially coding CAO were found in hybrid aspen T89 genome, Potrx053564g16785 (chlorophyll a oxygenase a; CAO1.1) and Potrx056793g18669 (CAO1.2). Potential sgRNAs for target genes were identified with CRISPR-P (http://cbi.hzau.edu.cn/cgi-bin/CRISPR2/CRISPR) including information regarding off target against other genome sequences in Populus genome. The potential off targets for sgRNAs were also checked against hybrid aspen T89 genome using BLAST method in plantgenie website (https://plantgenie.org/BLAST). sgRNA1 (GTGGAAGAAGGAGTTGCCACG) and sgRNA2 (GGATCCTCAACATCAAAGAG) were designed to target CAO1.1; sgRNA3 (GTAAGATGCCACAGTTTAAA) and sgRNA4 (GACCTTGGTTCAGTGAATGA) to target CAO1.1 and CAO1.2 together (mentioned here as CAO2). sgRNAs were introduced into entry vectors by site-directed mutagenesis PCR. GreenGate entry and destination vectors were acquired from Addgene. The final vector (containing promoter, Cas9 CDS, terminator, two sgRNAs and resistance cassette) was assembled by GreenGate reaction as described in André et al. (2022). *Escherichia coli* strain DH5α was used for amplification of all plasmids, which were then confirmed by sequencing (Eurofins). Vectors with different combinations of sgRNAs were transformed into Hybrid aspen T89 using a standard protocol. At least 30 individual transgenic lines from each transformation were screened for target gene deletions or SNPs using PCR and confirmed by sequencing. Forward: 5’-ACTTGCAAGCTAATCACACC-3’ and reverse: 5’- CAAGGTGAGACTTGAAAGTC-3’ primers were used to genotype transformation CAO1; forward: 5’-TGTGATCGGTGTGTGATTTTC-3’ and reverse: 5’- TATCACCAAATTTTCAATAC-3’ were used to genotype CAO1 gene in transformation CAO2; forward: 5’-ATAGTTTCCTGCTAAGGCTA-3’ and reverse: 5’-TCCCAGCCTGATGTTTATTG-3’ were used to genotype CAO1.2 in transformation CAO2. CRISPR of CAO1 transformation caused deletion lines, while CAO2 introduced stop coding in sgRNA3 location in both CAO1 and CAO2 genes producing a fragment of 67 amino acids instead of expected full CAO that formed from around 535 amino acids.

### Pigment extractions

Absorption spectra of leaves and thylakoids extractions in 80% acetone was recorded by UV-Vis spectrophotometer Shimadzu UV-2600i. Total amount of chlorophylls (a + b) and carotenoids were obtained by fitting the spectra as has been previously described by Croce et al. (2002). From these, ratios of chlorophyll a/b and chlorophyll/area were calculated.

### Thylakoid membrane isolation and biochemical analysis

Thylakoids were prepared as described by Caffarri et al. (2009), from overnight dark-adapted plants. BN-PAGE was performed on 4–12%-gradient (Järvi et al. 2011). 8 μg of chlorophylls were solubilized in a final detergent concentration of 2% b-DM and then loaded in each lane. 2D separation was performed from first dimension strips native-PAGE as described by Järvi et al. (2009). After electrophoresis, the proteins were visualized by Coomassie Blue staining.

### Integrated chlorophyll fluorescence by PAM fluorometry

Chlorophyll fluorescence was measured by a Dual PAM-100 (Walz). Plants were dark adapted for 1 before measurements. Saturating pulse (SP) of 10 000 μmol photons m^−2^ s^−1^ of 500 ms was applied to estimate the maximal fluorescence in the dark-adapted (F_m_) and light-adapted state (F’_m_). Induction of NPQ was achieved by an actinic light of 1000 μmol photons m^−2^ s^−1^. To record the NPQ kinetics, a SP was applied between 20 s for 8 min of actinic light. After, actinic light was turned off an SP was applied after 20 s, followed by 10 SP with a delay of 1 min. Intensity dosage per SP was 5000 μmol photons m^−2^ s^−1^.

We estimated NPQ from chlorophyll quenching fluorescence by the saturating light pulse method, where NPQ as a function of time, NPQ, can be described as:

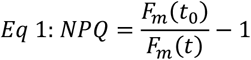

In the equation above, F_m_ (t_0_) is the yield of maximal fluorescence in a dark-adapted state, or F_m_, and F_m_ (t) is the maximal fluorescence measured at any time during the NPQ kinetic, also referred as F’_m_ (Baker, 2008). Eq. 1 is a simplified description of the integrated chlorophyll fluorescence intensity measured by PAM fluorometry, which can be formally expressed as an integral of fluorescence as a function of wavelength:

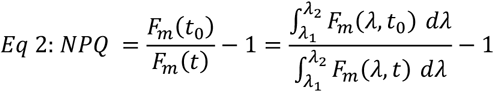

In Eq. 2, λ_1_ and λ_2_ are the wavelengths where the long-pass filter of PAM fluorometry integrates the fluorescence intensity. *In folio*, self-reabsorption can mislead spectral data interpretation. To avoid that, F_m_(t) can be corrected by the reflected and transmitted radiation using the correction factor “*f*” which is λ dependent (Chukhutsina et al., 2019):

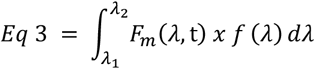

Where “*f*”:

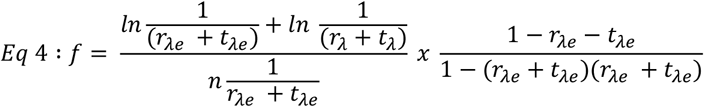

Here, r and t denote the reflected and transmitted radiation, respectively, while λe refers to the excitation wavelength and λ to the detection wavelength. As long as the self-reabsorption properties remain unchanged, for example during a NPQ induction using red-actinic light (Wilson & Ruban, 2020), NPQ corrected would be calculated as:

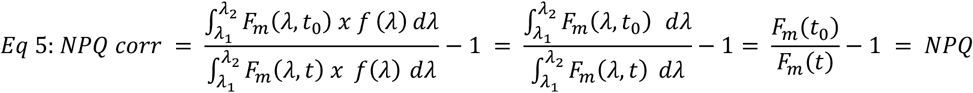

Therefore, any optical changes due to leaf structural changes would cancel out and NPQ will reflect only changes in PSII lifetime. This underlying assumption has allowed the photosynthetic community to study photosynthesis by PAM fluorometry (Schreiber et al., 1986) and explore from a vast collection of mutants the mechanisms that orchestrates NPQ. Note that the same assumptions are applied to the well-known parameters used to study energy partitioning or the redox state of the PSII RC (Baker, 2008), where:

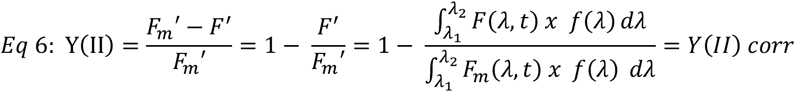

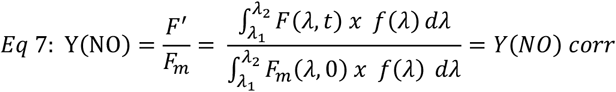

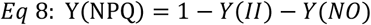

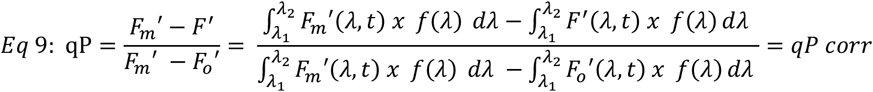

Here, F’ and Fo’ stand for the minimal yield of fluorescence at any given time during a NPQ kinetic and the minimal yield of fluorescence once all the reaction centers have been reopen, respectively. To determine Fo’ we used the approximation Fo’ = Fo/ (Fv/Fm + Fo/Fm’) as described by Oxborough and Baker (1997).

### Wavelength dependent NPQ inductions

Fast fluorescence inductions were recorded *in folio* from dark adapted leaves at 686, 700 and 730 nm using a ChloroSpec L1 (Nanda et al., 2024). Fo was set at 12.6 μs and the STF (single turnover flash) duration to 130 μs and an intensity of 60 000 μmol photons m^−2^ s^−1^. STF burst duration was 0.54 ms. A MTF (Multi-Turnover Flash) of 500 μs and an intensity of 15 000 photons μmol m^−2^ s^−1^ was applied subsequently. Values coming from the MTF were used to estimate the NPQ evolution spectra. To perform short NPQ kinetics, 1000 μmol photons m^−2^ s^−1^ were applied for a range of 8 min. A logarithmic sequence was applied to obtain MTF(λ,t), were each FI + MTF was spaced by 1, 1,1,1,2,3,6,12,23,30,44,65,100 and 191s. Subsequently, dark recovery was measured for 10 min applying a FI + MTF spaced by 5, 10, 20,40, 45, 60, 60, 60, 60, 60, 60, 60 and 60 s. Intensity dosage per FI + MTF was 1900 μmol photons m^−2^ s^−1^. Long NPQ kinetics were done at 1500 photons m^−2^ s^−1^ for 4572s followed of 1200 s of dark relaxation. Plants were dark adapted for 1 before measurements.

Here we studied NPQ spectral kinetics using red-actinic light and followed the changes of the maximal fluorescence emission spectrum over time, F_m_(λ,t) coming from the MTF. In this analysis, NPQ (λ,t) can be described as:

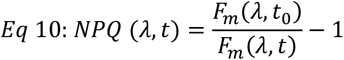

F_m_ (λ,t_0_) is the fluorescence emission spectra of a dark-adapted sample and F_m_ (λ,t) is the fluorescence emission spectra at any time during the NPQ kinetic. Note that Eq. 10 is identical to Eq. 1 and can be corrected by the correction factor “*f*” for self-reabsorption:

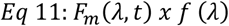

Therefore, it can be deduced that:

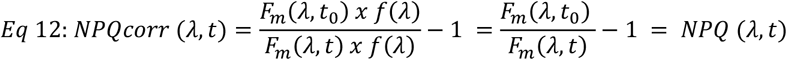

As above, any optical changes due to leaf structural changes would cancel out and NPQ (λ,t) will reflect only changes at PSII lifetime. As we induce NPQ by red-actinic light it is highly unlikely that self-reabsorption changes during these experiments hence spectra need not to be corrected, an assumption also made in PAM fluorometry. To follow the dynamics of new emitting species emergence during NPQ, we developed the parameter NESD.

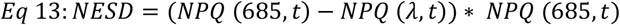

Here, the term NPQ (685,t) – NPQ (λ,t) captures the spectral heterogeneity of NPQ, thus the deviation of NPQ at any wavelength λ relative to the reference 685 nm, where PSII fluorescence dominates. Multiplying this difference by NPQ (685,t) incorporates the overall magnitude of NPQ, combining the heterogeneity by the magnitude of PSII-specific quenching.

Y(II) (λ,t), Y(NO) (λ,t) and Y(NPQ) (λ,t) was calculated according to Eq. 6, Eq. 7 and Eq. 8, respectively from each fast fluorescence induction detected at 686, 700 and 730 nm. From them, PSII energy partitioning heterogeneity was determined for Y(II), Y(NO) and Y(NPQ) by the following equations:

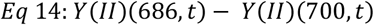

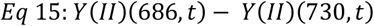

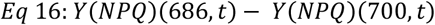

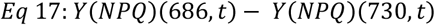

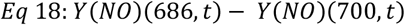

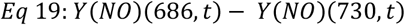

From these differences, dimensional reduction was applied by clustering analysis in combination with PCA.

### Integrated chlorophyll fluorescence and gas exchange

Net CO_2_ assimilation (A), stomatal conductance (gsw), intercellular CO_2_ (Ci) and chlorophyll a fluorescence were measured using an infrared gas analyzer (model LI-6800, LI-COR, Lincoln NE, USA) at 1500 μmol photons m^−2^ s^−1^, 21% O_2_ and 400 ppm CO_2_. During measurements leaf temperature was maintained at 25°C and humidity at 60%. Same conditions were applied to all measurements performed. Log were performed every 2 minutes by the auto-gen loop function in a light interval of 4680 s followed by a dark-interval of at least 1200 s. Each kinetic has a duration of at least 1:50 hs. A pre-dark phase of 240s before each induction was done to stabilize the samples and measure Fm. NPQ, Y(II), Y(NO), and qP were determined as mentioned above

### Statistical analysis

Statistical analysis was performed in Rstudio trough an in-house pipeline to analyze the raw data signals from ChloroSpec and processed data from Dual-PAM-100 and Licor-6800. To perform and plot the results from linear regression, PCA, cluster analysis, ANOVA and Tukey-test libraries ggplot2, stats, cluster, factoextra, multcomp and multcompview were used. For clustering supervised analysis was performed, where the optimal number of clusters was determined based on the elbow method.

## Results

### *Chlorina* mutants have a reduced physical PSII antenna size and NPQ

To study how LHCs and major regulators affect NPQ and the development of NES in photoinhibitory and non-photoinhibitory conditions in *Poplar*, we generated a collection of aspen *chlorina* mutants using CRISPR-CAS9, in combination with PsbS (*psbs*) and Violaxanthin de-epoxidase (*vde*) mutants (Nanda et al., 2025; Fig. 1). We identified two copies of the CAO gene, targeted one (*cao1*) or both (*cao2*) to partially or completely disrupt the synthesis of chlorophyll b (Fig. 1, Supplementary Fig. S1). This resulted in a set of aspen lines with varying CAO activity and, consequently, different chlorophyll b content. We found that an increase in chlorophyll a/b ratio was associated with a decrease in the chlorophyll content per unit leaf area (Fig.1 B-C). In *psbs* and *vde*, chlorophyll a/b ratios and chlorophyll content were the same as in wild type (T89; Fig. 1 B-C). As expected, changes in chlorophyll b content in *chlorinas* were reflected in a reduction of the PSII physical antenna size (Fig. 1 D-F, Supplementary Fig. S1 and S2). The lines *cao1.3* and *cao1.11* showed only small changes in their chlorophyll a/b ratio and thylakoid organization, whereas the absence of chlorophyll b (*cao2.14* and *cao2.18*) compromise LHCB accumulation and PSII supercomplex stability.

**Figure 1.**
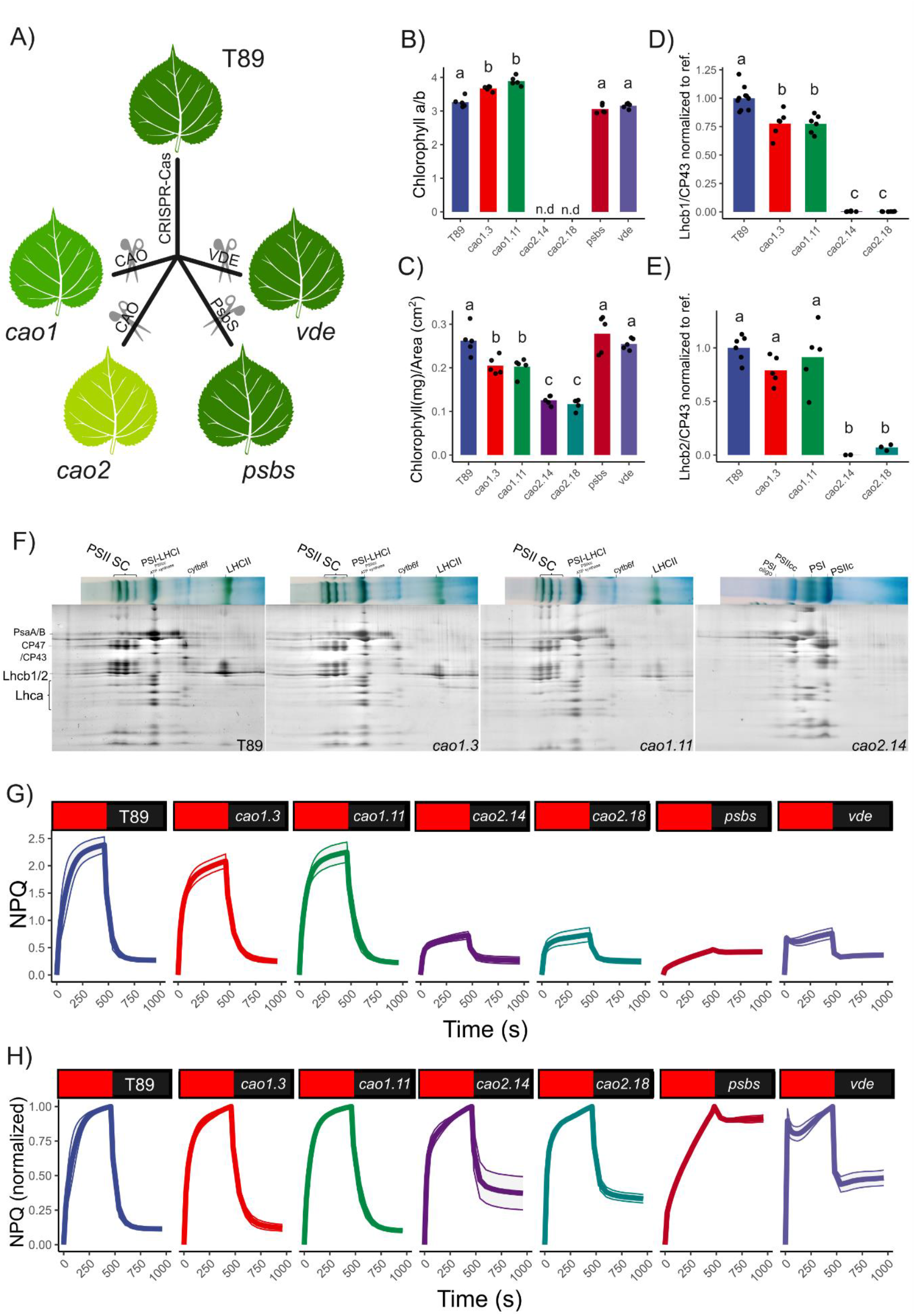
Lhcb, PsbS and zeaxanthin modulate NPQ in aspens. A) Schematic describing chlorophyll a oxygenase (CAO), PsbS and VDE mutants production by CRISPR-Cas9 in aspen (T89). B-C) Chlorophyll a/b and chlorophyll content per leaf from T89, *cao1, cao2, psbs* and *vde* mutants. Pigment content values are means from 5 biologically independent experiments; each point is from one replicate. Shared letters indicate non-significant differences between the groups (Tukey’s tests, p <0.05). Note that *chlorina* mutants *cao2.14* and *cao2.18* lack of chlorophyll b (n.d). D-E) Relative signal of Lhcb1 and Lhcb2 to CP43 was quantified by immunoblotting and normalized to T89. Shared letters indicate non-significant differences between the groups (Tukey’s tests, p <0.05). F) 2-D BN/SDS-PAGE from aspen thylakoids solubilized in 2% β-DM. 8 μg of chlorophylls were loaded per lane. In Supplementary Fig. S1 and S2, BN-Page and blots are shown. G) NPQ inductions performed at 1000 μmol photons m^-2^s^-1^ in collection of NPQ aspen mutants using a standard pulse amplitude modulated fluorometer (PAM). H) Normalized NPQ kinetics. On top of each trace is indicated the corresponding genotype. Induction and relaxation phase are represented by the red and black boxes, respectively. Data are mean ± s.d (n = 5 biologically independent experiments).

Figure 1 illustrate that NPQ in aspen is PsbS, Zea and LHCB dependent. *Cao1* show NPQ inductions patterns like T89 but NPQ is strongly reduced in *cao2* lines, as expected for mutants lacking LHCII (Fig. 1 G-H). Additionally, both *psbs* and *vde* show a marked reduction in NPQ reflecting the absence of qE and qZ components, respectively. When NPQ was normalized (Fig. 1 H), we observed the rapid induction of the qE-associated component in T89, *cao1, cao2*, and *vde*, but not in *psbs*, as expected.

### NPQ evolution spectra reflects changes in PSII lifetime *in folio*

A limitation of PAM fluorometry is that the fluorescence signal is integrated above a certain detection cutoff wavelength, hence potential variations in the short-wave emission spectrum where PSII contribution dominates are ignored. To break the bidimensional limit of NPQ, it is necessary to time-resolve the fluorescence emission spectra during light inductions (Nanda et al., 2024). We performed NPQ spectral kinetics using red-actinic light and monitor the changes of the maximal fluorescence emission spectrum over time, Fm (λ,t). We derivate NPQ (λ,t) as described in Eq. 10, where self-reabsorption cancels out, as happens for PAM fluorometry (Eq. 5). Therefore, NPQ (λ,t) and NPQ reflects changes in PSII fluorescence lifetime at any given time.

We found that NPQ (λ,t) depends on PsbS, Zea and on the LHC antenna (Fig. 2, Supplementary Fig. S3, S4, S5 and S6). In T89 and *cao1*, NPQ spectra exhibit a canonical behavior like other higher plants (Nanda et al., 2024). However, the differences in the NPQ spectra were absent in *cao2, psbs* and *vde*. To quantitatively estimate NPQ (λ,t) changes we introduce a new coefficient named NES development (NESD), described in Eq. 13. In this specific conditions, we compare NPQ (685,t) with NPQ (720,t) where the largest spectral differences are found for the NPQ spectra of T89 (Fig. 2 C).

**Figure 2.**
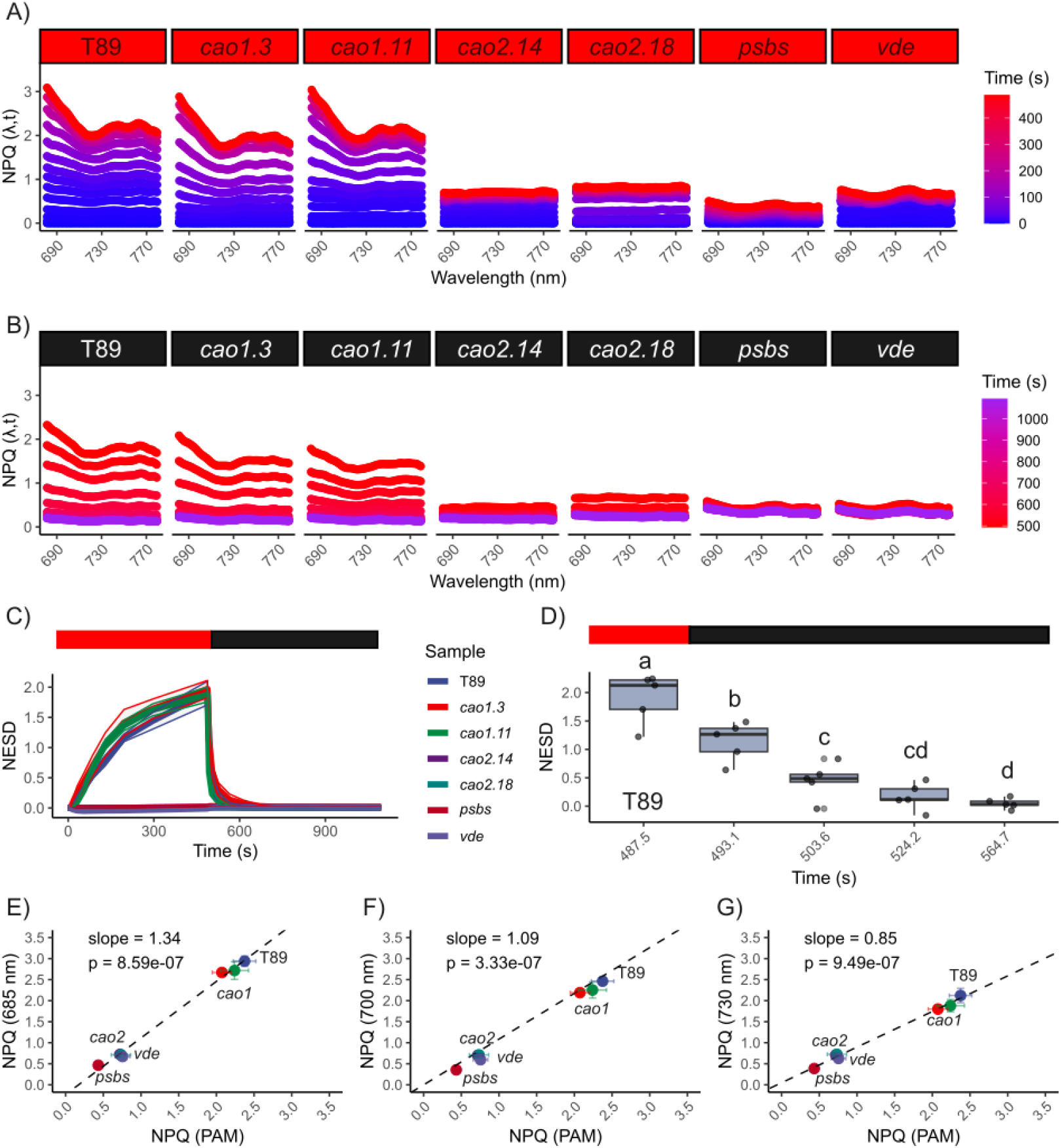
Lhcb, PsbS and zeaxanthin are required for the development of new emitting species (NES) *in folio* under non-photoinhibitory conditions. NPQ inductions were performed at 1000 μmol photons m^-2^s^-1^ in collection of NPQ aspen mutants. Induction and relaxation phase NPQ evolution spectra are shown in A) and B), respectively. Each NPQ evolution spectra represents one biological replica. C) New emitting species development (NESD) was calculated from NPQ evolution spectra as (NPQ_(685,t)_ – NPQ_(λ,t)_) * NPQ_(685,t)_. Data are mean ± s.d (n = 5 biologically independent experiments). D) NESD relaxation in T89. Shared letters indicate non-significant differences between the groups (Tukey’s tests, p <0.05). Linear regression was performed from NPQ ± s.d detected at different λ and compared to the NPQ ± s.d from Figure 2 obtained from integrated PAM fluorometry, NPQ (PAM). In E), F) and G) regression is shown for 685, 700 and 730 nm, respectively. Slope and associated p-values is indicated for each regression.

In T89 and *cao1* the differences between NPQ (685,t) and NPQ (720,t) were pronounced, noticing that NESD follows the dynamics of classic NPQ inductions and relaxations. In contrast, we found that NESD tends to 0, in *cao2, psbs* and *vde*. Thus, during NPQ, the presence of LHCB, PsbS and Zea is necessary for the development of NES. Moreover, NESD is highly dynamic and after its induction at the light phase, is relaxed up to 75% in 16 s and abolished after 40 s. (Fig. 2 D). Compared to “standard” NPQ measurements using PAM, NPQ (λ,t) in *npq* and *chlorina* lines could be up to 30% higher at shorter detection wavelengths indicating that PAM fluorometry underestimates NPQ (Fig. 2 E-G).

### PSII energy partitioning heterogeneity emerges from the interaction of the leaf light gradient and NPQ regulation

PSII energy partitioning is a valuable tool for understanding the fate of excitons in photosynthesis. The pioneering work of Genty et al. (1989) directly linked chlorophyll a fluorescence to photosynthesis, introducing the parameter Y(II) and later Y(NO) and Y(NPQ) were derived (Baker, 2008; Klughammer & Schreiber, 2008). These parameters are typically measured under red-actinic light conditions, where self-reabsorption effect cancel out (see above, Eq: 6-8) therefore they do not need to be corrected for reabsorption in PAM fluorometry analyses, nor using our wavelength-dependent fluorescence method.

In Supplementary Figure 7, results from PAM fluorometry are shown. Upon NPQ inductions (1000 μmol photons m^-2^s^-1^; Fig. 1) once the system reaches steady state in T89, Y(NPQ) dominates followed by Y(NO) which constrains the use of energy for photosynthesis by PSII, known as Y(II), where only 5∼6% of absorbed photons are used for charge separation. In line, *cao1* show a similar trend. However, under the absence of qE or LHCII, the system becomes mainly dominated by Y(NO), which can be up to two times larger than T89.

Supplementary Figure 8 display traces of Y(II), Y(NPQ), and Y(NO) calculated from fluorescence induction detected at three different wavelengths. As these variables showed a complex behavior, we next focused our analysis on the final point of the light phase induction (Fig. 3). In T89, Y(II) typically decreased at shorter detection wavelengths while Y(NPQ) increased. As monochromatic red actinic light causes most excitation at the top of the leaf, differential excitation could be accompanied by a decreasing ΔpH and NPQ gradient as the light penetrates the leaf deeper. Thus, NPQ gradient could affect PSII energy partitioning at different wavelengths (Fig. 3 A).

**Figure 3.**
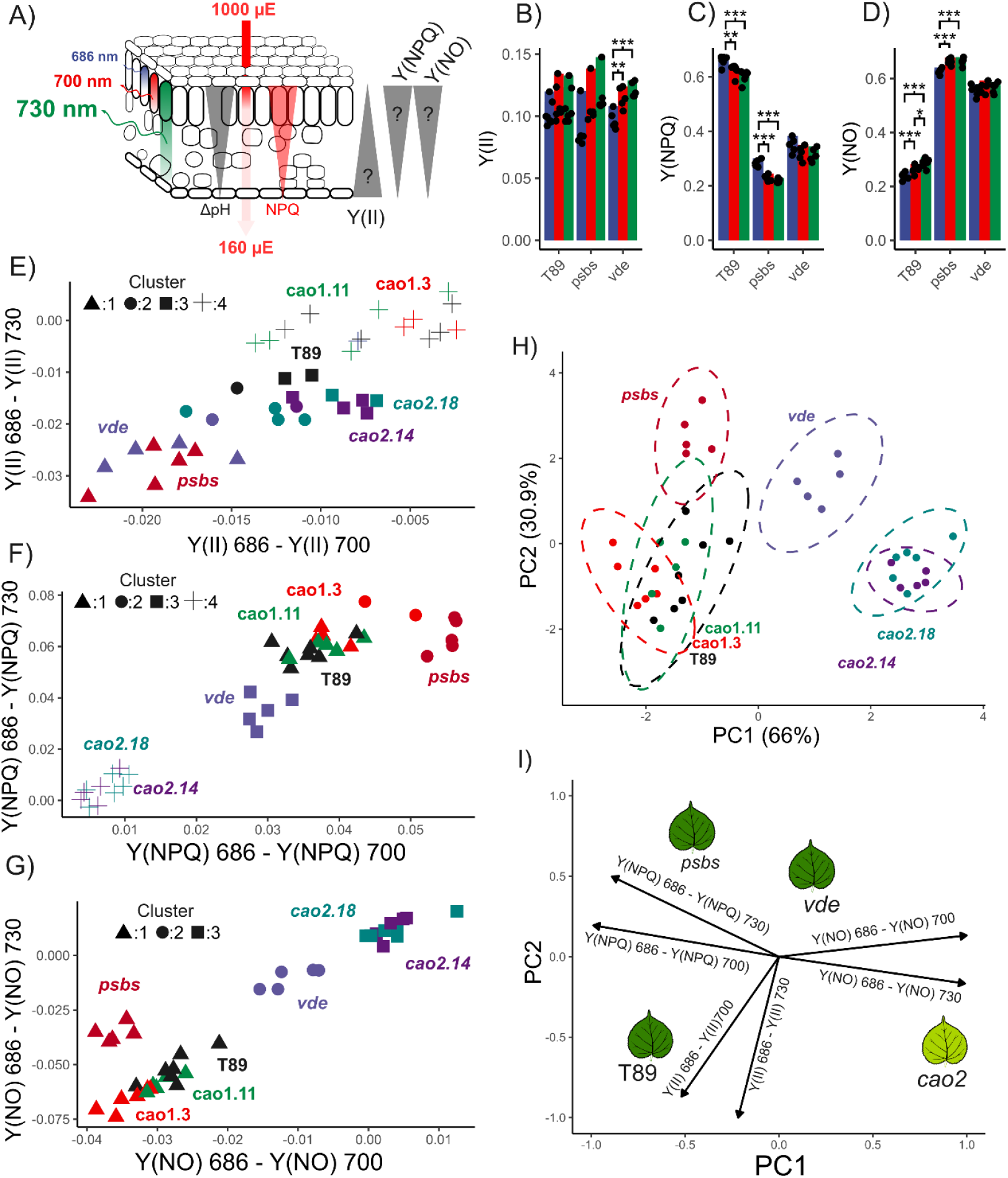
PSII energy partitioning heterogeneity upon NPQ is mutant dependent. A) A schematic illustrating the relationships between monochromatic excitation through the leaf blade, known as light gradient, (depicted by the red arrow) and the resulting NPQ/ΔpH changes, represented by red and grey triangles, respectively. It is expected that, under monochromatic excitation, PSII excitation will be greater on the abaxial side of the leaf compared to the adaxial side, leading to larger stromal-luminal ΔpH and NPQ. In this case, it is anticipated that Y(II), Y(NPQ), and Y(NO) will follow the NPQ gradient. B), C), D) ANOVA statistical analysis was applied to Y(II), Y(NPQ), and Y(NO), respectively, derived from NPQ inductions at 480s, when the system reached steady state in the T89 genotype. Significant differences are indicated by *, **, and ***, with p-values lower than 0.05, 0.01, and 0.001, respectively. E), F), G) Multidimensional reduction of the data shows differences in Y(II), Y(NPQ), and Y(NO) measured at 686 nm and compared to 700 nm, and 730 nm for each biological replicate. Clustering analysis was performed using the k-means method, with the number of clusters per condition determined by a supervised analysis using the elbow method. Each point represents a distinct biological replicate (n > 5). H) Principal component analysis (PCA) was performed on the differences of Y(II), Y(NO), and Y(NPQ) recorded at 686 nm to 700 nm and 730 nm. Six variables were computed for each biological replicate. PC1 accounts for 66% of the variation, which is primarily influenced by the differences in Y(NPQ) and Y(NO), as described in the loading plot (I). PC2 explains 30.9% of the variation, mainly driven by differences in Y(II). In panel H, each ellipse represents a 95% confidence interval for each genotype based on its centroid.

To investigate this, we performed an ANOVA on Y(II), Y(NPQ), and Y(NO) at different detection wavelengths for T89, *psbs*, and *vde* (Fig. 3 B-D). Y(II) did not show significant variation across different wavelengths for T89 and *psbs* but was reduced in *vde* at shorter wavelengths. In contrast, Y(NPQ) increases at shorter detection wavelengths for T89 and *psbs*, whereas the opposite effect was observed for Y(NO). In *vde*, no significant differences were found for Y(NPQ) and Y(NO) at the different detected wavelengths. However, this approach may underestimate the spectral differences between each biological sample, as genotype-specific variations were masked by overall differences within each group and a complete analysis of this dataset for a single time point required of 63 comparisons (Supplementary Fig. S9 and S10). Accordingly, we designed a novel pipeline applying dimensional reduction.

Dimensional reduction is a common technique used in data analysis that simplifies complex datasets by reducing the number of variables while preserving essential information (Van der Maaten et al., 2009). This approach is valuable when dealing with high-dimensional data, such as that generated by simultaneous detection of fluorescence, where each sample has a heterogeneous spectral signature for Y(II), Y(NO) and Y(NPQ) which can be defined by Eq. 14-19. From these differences we performed cluster analysis comparing Y(II), Y(NO), and Y(NPQ) at 686 nm with 700 nm, and 730 nm at the end of the NPQ induction (Fig. 3 E-G). When the cluster analysis was combined with principal component analysis (PCA) to examine the sources of heterogeneity in PSII energy partitioning (Fig. 3 H-I) 66% of the variation in energy partitioning heterogeneity (PC1) was explained by the antagonistic relationship between Y(NPQ) and Y(NO), while 31% (PC2) was explained by Y(II).

T89, *cao1.3*, and *cao1.11* clustered together in the PCA and cluster analysis, suggesting that a moderate reduction in chlorophyll and LHCII content does not significantly affect how plants cope with increasing light intensities. As expected, these lines were primarily affected in Y(NPQ), likely associated with the leaf light gradient. Here, Y(NPQ) showed an antagonist relationship with Y(NO); at 686 nm larger values of Y(NPQ) were associated with lower Y(NO) values, whereas at longer wavelengths, the opposite trend was observed. (Fig. 3 F-G). Surprisingly, *psbs* and *vde* had a larger effect on PSII energy partitioning heterogeneity, despite lacking NESD and qE. In *psbs*, Y(NPQ) can be up to 23% larger at 686 nm than 730 nm (Supplementary Fig. 10), exhibiting the most pronounced effect in Y(NPQ). This is accompanied by its antagonistic relationship with Y(NO), like T89 response, a pattern supported by the PCA where their ellipses slightly overlap (Fig. 3 H). In *vde*, the strongest effect was observed in Y(II), while the typically antagonistic interaction between Y(NO) and Y(NPQ) appeared to be attenuated. Likewise, the absence of LHCII and the strong reduction in chlorophyll content in *cao2* had only a significant effect on Y(II) heterogeneity (Fig. 3 I). The cluster analysis indicated that *cao2* exhibited the least heterogeneity in Y(NPQ) and Y(NO), similarly to the effect observed in *vde* suggesting that some properties might be shared between both lines.

### Sustained quenching could be uncoupled from photoinhibition

PCA revealed differences in energy partitioning among the *npq* mutants, with *vde* and *cao2* showing the largest divergence from T89 (Fig. 3). Both lines show a weak antagonist effect between Y(NPQ) and Y(NO) heterogeneity suggesting that they might be more susceptible to photoinhibition, as is expected from mutants lacking qZ or LHCB proteins (Nilkens et al., 2010; Havaux and Tardy, 1997). To study the relationships between PSII energy partitioning heterogeneity, NESD, photoinhibition and sustained quenching we performed a long NPQ induction experiment in highlight (HL)/photoinhibitory conditions (80 minutes in 1500 μmol photons m^−2^ s^−1^, 400 ppm CO_2,_ 25°C and RH 60%), followed by relaxation for 20 minutes in our aspen mutants (Fig. 4), combining measurements of CO_2_ assimilation and PAM fluorometry (Li-6800; Li-COR). To get a numerical value of the amount of quenching that was sustained/slowly relaxing in this experiment, we used the equation SQ = 1 – (NPQ_max_ – NPQ_t_)/NPQ_max_), which equates to NPQ normalized. In parallel we monitored NESD by spectrally resolved fluorescence inductions. Although gas exchange measurements typically are performed using a combination of blue and red light, we used here only red-actinic light to allow for comparisons and avoid changes in self-reabsorption during the kinetics.

**Figure 4.**
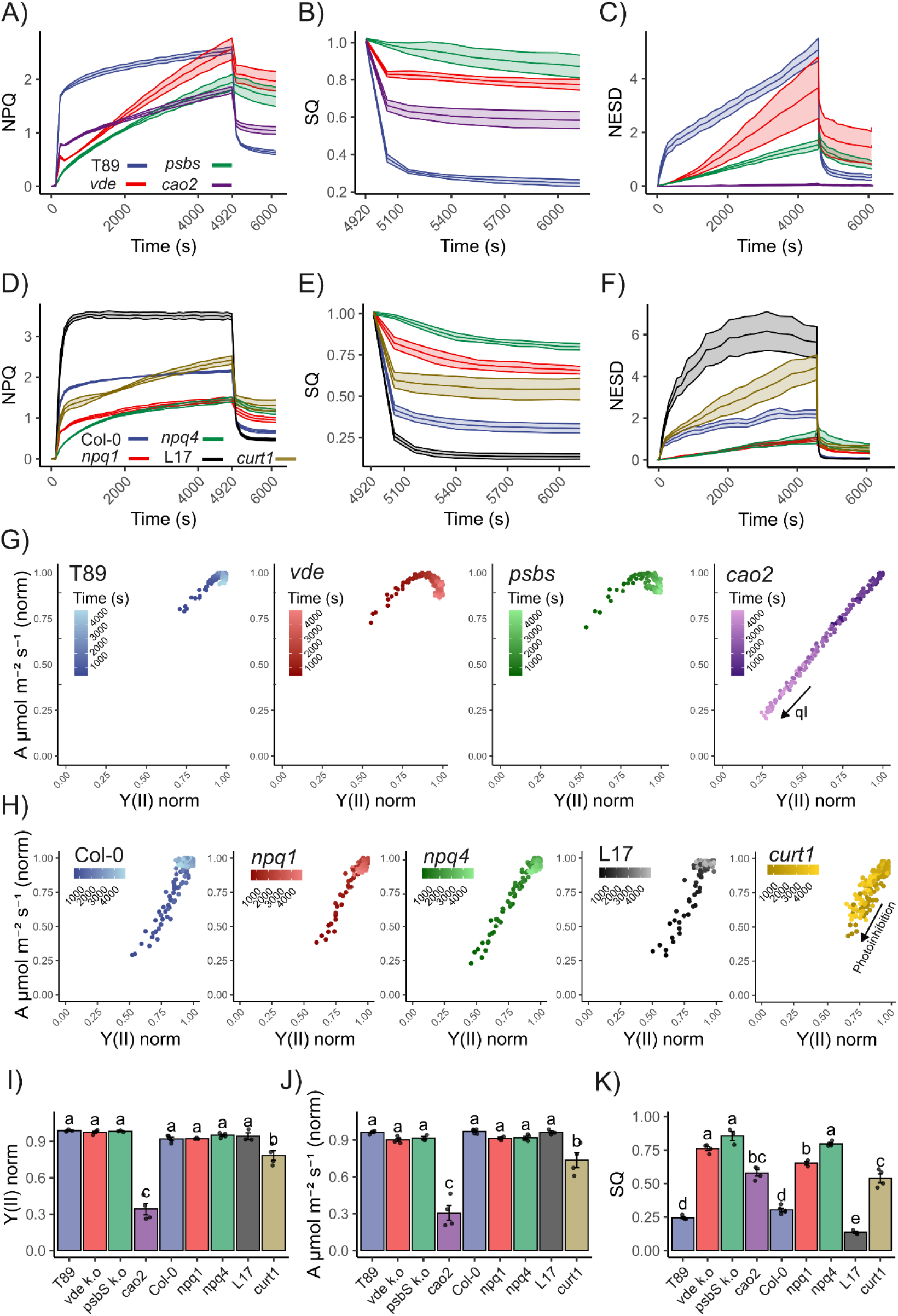
Relationship between NESD, photoinhibition and sustained quenching in aspen and *Arabidopsis* mutants. A-D) Aspens and Arabidopsis NPQ inductions by integrated fluorometry (Li-COR) when illuminated with 1500 μmol photons m^-2^s^-1^ for 4680s followed of a relaxation phase of 1200s. Before the light phase a 240s dark interval was recorded to stabilize the sample and determine Fm. B-E) Sustained quenching (SQ) was determined as 1 – (NPQ_max_– NPQ_t_)/NPQ_max_ from A) and D), respectively. C-F) NES development was obtained from spectrally time-resolved kinetic in similar conditions. For L17, NESD was scaled to 0.5x. G) Gas exchange and fluorescence was monitored at 400 ppm and 1500 μmol photons m^-2^s^-1^ for 4680 s. Each biological replica was normalized to the maximal value of Y(II) and CO_2_ assimilation after full activation of NADPH reductase (≥ 480 s). The evolution of CO_2_ assimilation as a function of Y(II) is presented during the light induction phase, where time progress from dark (480 s) to light colours (4920 s). H) Gas exchange and fluorescence was monitored as described in C) for *Arabidopsis* mutants. Data is mean ± s.e of at least 3 independent biological replicas. I-J) Normalized Y(II) and CO_2_ assimilation at actinic light end point (4920 s.). K) SQ after 20 minutes of dark relaxation (6120 s). Shared letters indicate non-significant differences between the groups (Tukey’s tests, p <0.05). Each point represents a distinct biological replicate (n > 3).

In this HL regime, T89, *psbs* and *vde* kept up CO_2_ assimilation throughout the experiment (*psbs* and *vde* somewhat lower than T89) while it decreased much over time in *cao2* (Fig. 4 J, Supplementary Fig. S13). During this treatment also *psbs* and *vde* developed significant quenching (Fig. 4 A), at the end of the induction phase *vde* had even higher NPQ than T89. This quenching relaxed slowly, NPQ decreased only with 20-25 % in *psbs* and *vde* compared to 75 % in T89, while cao2 showed a 40% decrease (Fig. 4 B and K). Under these conditions, *psbs* and *vde* also developed NES (Fig. 4 C, Supplementary Fig. 11) albeit significantly less than T89, whereas cao2 did not develop NES. If normalized CO_2_ assimilation is plotted against normalized Y(II), CO_2_ assimilation in *cao2* declined almost proportionally to Y(II) down to ca 25 %, while both parameters instead increased in T89, *psbs* and *vde* (Fig. 4 G). Similar effects were observed when normalized qP was compared to A (Supplementary Fig. 15). Hence, *cao2* suffered from photoinhibition during these conditions without developing NES while *psbs* and *vde* developed sustained quenching and NES with photoinhibition comparable to T89 (Fig. 4 J). Although, no changes were observed in stomatal conductance (gsw) for all the lines analyzed, the intracellular CO_2_ (Ci) increase up to 25% in *cao2*, suggesting and independent regulation of stomatal closure by CO_2_ signaling in *cao2* (Engineer et al., 2016)

To confirm the unexpected results of *psbs* and *vde* in another angiosperm species, we conducted a long-term NPQ kinetics (same conditions as described above) experiment using *Arabidopsis* lacking PsbS (*npq4*) or VDE (*npq1*), or overexpressing PsbS (L17). To see if thylakoid structure was important for these traits, we also included the quadruple mutant of CURVATURE THYLAKOID 1, *curt1*, strongly affected in thylakoid stacking and plasticity. The results were similar, CO_2_ assimilation was kept up but quenching was induced also in *npq4* and *npq1* after extended exposure to HL and, like in aspen, the quenching was only slowly relaxing (Fig. 4 D, E and H-K Supplementary Fig. S14). NES were developed in all genotypes, where L17 reach values up to 12 and large sustained values in the relaxation phase (Fig. 4 F, Supplementary Fig. S12 and S16). Finally, *curt1* had a strong NESD and significant sustained quenching, but up to 50% decrease in CO_2_ assimilation and in Y(II), as well as for qP (Fig. 4 E-K, Supplementary Fig. S15). In these conditions, a non-significant decrease was observed in gsw for L17 whereas curt1 showed a 50% decline compared to Col-0, indicating that the decline in CO_2_ in *curt1* could be linked to the redox state (Wang et al., 2016). Accordingly, *curt1* showed a decline of 30% in the amount of open reaction centers (qP) when compared to Col-0 at the end of the high light treatment. Lastly, when analyzing Y(NO), we found that *cao2* exhibited a 150% increase during the relaxation phase compared to its reference value in dark-adapted state, while *curt1* showed similar values (Supplementary Fig. S13 and S14).

The differences in these traits are summarized for the mutants lacking different proteins under standard conditions in Figure 5 A, and under photoinhibitory conditions in Figure 5 B.

**Figure 5.**
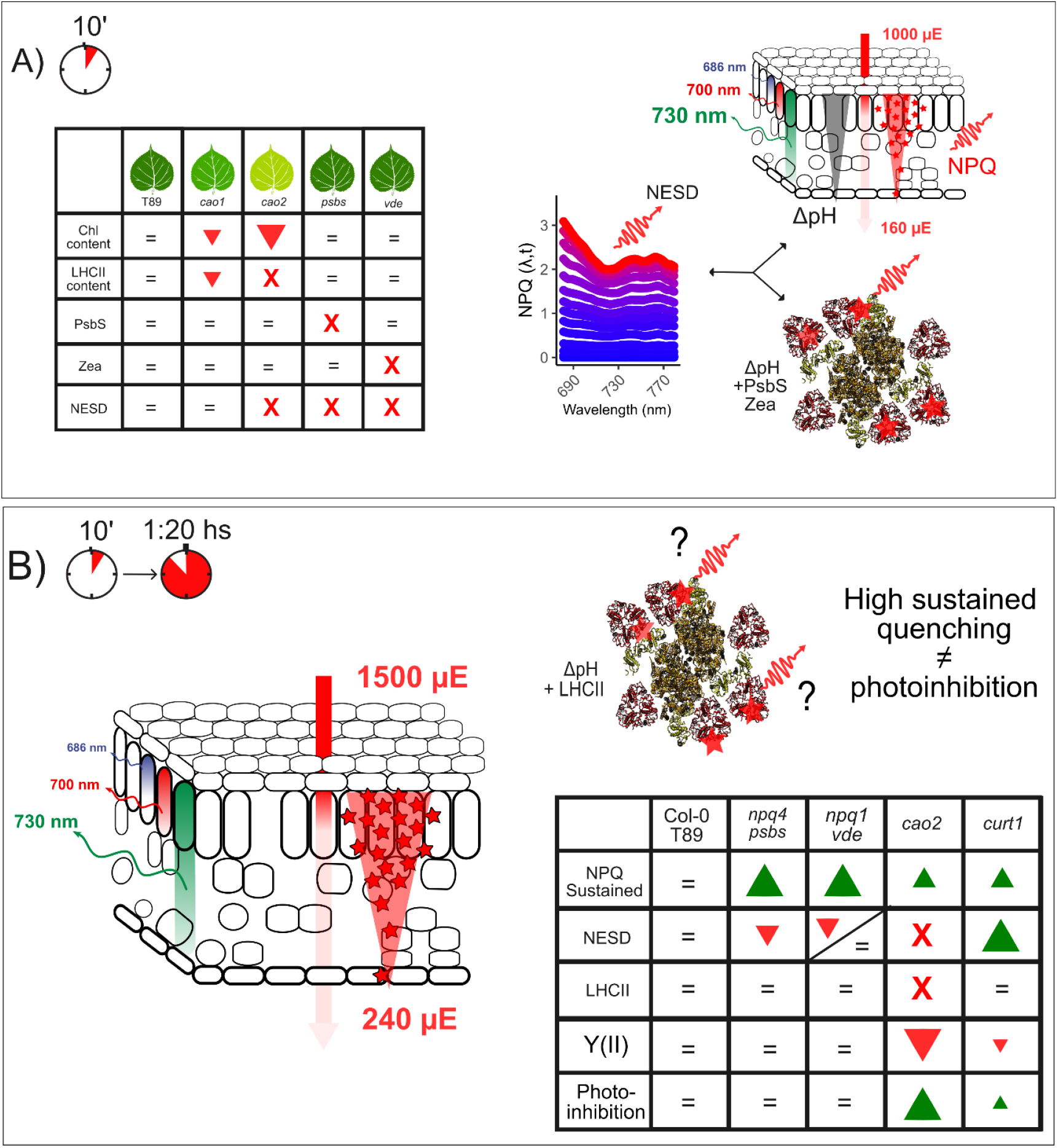
The relations between NESD, photoinhibition and sustained quenching. A) Table summarizing the relationships between chlorophyll content (Chl), LHCII content, PsbS, zeaxanthin formation (Zea), and the development of new emitting species (NESD) in response to fast NPQ inductions within the leaf. The red triangles and their respective sizes indicate the direction and magnitude (decrease) of phenotypic changes relative to the T89 genotype. Red crosses denote the absence of the specified variable. A schematic illustrating the relationships between NPQ gradient (represented by red triangle), light gradient (depicted by a red arrow), and the quenching activities of Zea, PsbS, and LHCII at photosystem II (red stars). The interplay of these factors leads to a novel property of NPQ characterized by a significant reduction of NPQ above 700 nm, associated with the emergence of NES. B) Under sustained quenching the development of NESD is independent of PsbS and Zea, where the only pre-requisite is the presence of Lhcb proteins. A table summarizing the relationships between sustained quenching (NPQ sustained), NESD, LHCII, Y(II) and photoinhibition is presented. The red triangles indicate decrease compared to their reference line whereas the green triangles increase.

## Discussion

### What produce new emitting species during NPQ?

In recent years, the study of photoprotective sustained quenching and photoinhibition has gained a lot of attention (Malnoë et al., 2018; Bag et al., 2020; Nawrocki et al., 2022; Bru et al., 2022; Smith et al., 2024). Although, these processes are important for ecosystems and agricultural productivity they are quite poorly understood and its definitions from time-course fluorescence data after certain treatments, are not straightforward. Malnöe (2018) argues that “qI has rightfully been called “ill-defined” (Nilkens et al., 2010) and the source of confusion in the field comes from the term “photoinhibition” as (Maxwell and Johnson, 2000) explained: “it is important to note that this term, when applied to fluorescence analysis, generally refers to both protective processes and to damage to the reaction centers of PSII (Osmond, 1994), whilst in more molecular studies, it is specifically the latter that is referred to as photoinhibition”. If we regard qI-photoinhibition as quenching at the core of PSII (Nawrocki et al., 2022), qI occurs when the photoprotective machinery have failed to protect PSII and represents a costly mechanism, negatively affecting oxygen evolution and productivity. Sustained quenching is part of mechanisms protecting the PSII RC to avoid photodamage and therefore could prevent the development of qI. In plants, the antenna has a pivotal role balancing light harvesting and quenching and one can assume that NPQ, including sustained quenching, involves LHCB complexes (Ruban and Saccon, 2022; Bru et al., 2022). We have also reported one sustained photoprotective quenching mechanism in overwintering gymnosperms, PSII protection by direct energy transfer to PSI (Bag et al., 2020), demonstrating the mechanistic diversity of sustained quenching across plant lineages to different environmental cues.

By using spectrally resolved fluorescence and net photochemical productivity, monitored by gas exchange analysis, we demonstrate when qI is coupled or not to sustained quenching, and the spectral signatures associated to these processes. In increasing light, and in the absence of photoinhibition, plants develop so-called NES (Fig. 3), and two models have tried to explain their origin. The idea that NES are linked to qE/PsbS/LHCII (see for example Horton et al. 2005; Holzwarth et al. 2009; Johnson and Ruban, 2009) has been challenged by recent studies, suggesting that their origin is product of closed PSII RC under actinic light (Farooq et al. 2018). Here we explored in the absence of photoinhibition how the decrease of PSII absorption cross section and the absence of NPQ modulators affect the development of NPQ spectra, focusing on poplar— a model fast-growing tree in which light harvesting remains largely unexplored. Upon NPQ induction, T89 show a classical NPQ evolution spectrum, where we observed NES at longer wavelengths (Fig. 2). We also show that when comparing ref. lines and *npq* mutants, differences in the emission spectra lead to an underestimation of up to 30% in NPQ, when measured by PAM fluorometry. Recently, it has been reported NPQ values close to 3 at 680 nm in *Arabidopsis* leaves by chlorophyll fluorescence lifetime snapshots (Lam et al., 2024), in agreement with our previous report where larger differences are observed in the NPQ evolution spectrum (Nanda et al., 2024). Obviously, NES could lead to a significant underestimation of NPQ using “standard methods”.

We suggest a new parameter – NESD - that can be calculated from this type of data to obtain time-resolution of the spectral differences of NPQ. This parameter can reflect the dynamics of the process, an aspect that cannot be monitored using ultrafast time-resolved spectroscopy which measure the lifetime of chlorophyll fluorescence during steady-state conditions (see e g Chukhutsina et al. 2019). In native systems, NES are rapidly both induced and relaxed, with kinetics like qE development (Fig. 2). Such a fast relaxation would not make comparable the PSII emission spectra in the closed state during NPQ and the open state during the relaxation phase, like in Farooq et al. (2018). Moreover, we found no NESD in c*ao2* mutants lacking antenna complexes; if closed PSII RC were the source of NES, then cao2 would be expected to show heterogeneous NPQ spectral profiles over time. Instead, their spectra remain uniform (Fig. 2 and 4). Taken together, these observations support the conclusion that, under NPQ conditions, is unlikely that closed PSII RCs are the source of NES. Under long NPQ inductions at 1500 μE, where significant sustained quenching was established in T89, NESD could only be detected after 1 minute of dark relaxation, was nearly absent in the Col-0 line but enhanced in L17 overexpressing PsbS (Fig. 4, Supplementary Fig. S16). This suggests that the findings from Farooq et al. (2018) might be due to more complex emerging properties.

### Light gradient, qE and PSII energy partitioning heterogeneity – a complex relationship

NPQ spectra and NESD in the leaf seem to be connected to PsbS, Zea, NPQ and photoprotection but how? We think we could rule out the possibility that NPQ spectra and NES is affected by self-absorption in the leaf, based on the arguments around Eq. 12 above. However, NES might develop as a consequence of leaf differential excitation, known as light gradient (Fig. 3 A). When light strikes the upper surface of a leaf, it is not distributed evenly across all tissue layers. Photosystems located in the upper layers of the leaf absorb more light and become more excited than those in the deeper layers. Thus, the light gradient produces a NPQ gradient across the leaf. As a result, NESD may manifest as a gradual decline in qE with depth, which cannot be corrected using Eq. 3.

However, if the light gradient were the sole explanation for NESD, *psbs, vde* and T89 – despite having similar chlorophyll content – would be expected to exhibit comparable NESD. Moreover, T89 and *cao1* display comparable NESD, although *cao1* has a reduction in chlorophyll content, which would typically produce a more uniform light gradient. Instead, our data suggests that NPQ and NESD should be considered emergent properties resulting from the interplay between the light gradient and changes in thylakoid re-organization which together influence PSII fluorescence lifetimes during NPQ and its apparent spectra. Together, these emergent properties modulate the fate of energy utilization at different wavelengths, i.e. PSII energy partitioning heterogeneity (Fig. 3).

Under our experimental conditions, greater discrepancies in PSII energy partitioning heterogeneity were observed among T89, *psbs, cao2*, and *vde*. The light gradient appears to create an antagonistic relationship between Y(NPQ) and Y(NO), while Y(II) remains relatively unaffected in T89 (Fig. 3). This suggests that Y(II) is evenly distributed across different chloroplast layers, maintaining consistent quantum efficiency performance. Additionally, the higher Y(NPQ) and lower Y(NO) values observed at 686 nm indicate an appropriate response to the increased excitonic pressure at the upper surface of the leaf. As expected, a similar response was seen in *cao1*, and surprisingly, also in *psbs*, indicating that these mutants effectively respond to the light gradient. Interestingly, PAM fluorometry reveals that *psbs* is predominantly influenced by Y(NO), although this effect is mainly observed at longer detection wavelengths (Supplementary Fig. S8 - S10). However, *vde* and *cao2* exhibited a different pattern, where primarily Y(II) was influenced, and the antagonistic relationship between Y(NPQ) and Y(NO) heterogeneity was largely reduced, although *vde* has a similar chlorophyll content than T89. Overall, these findings highlight the importance of simultaneous chlorophyll fluorescence detection to gain a deeper understanding of PSII energy partitioning in plants, a process influenced by the interaction between the light gradient and NPQ.

### When sustained quenching uncouples from photoinhibition

Lack of qE will increase excitonic pressure on PSII and influence the RC state, increasing the formation of a slow NPQ component named δ by Ramakers et al. (2024), assumed to represent qI or a qI-like process (Li et al., 2002; Nilkens et al., 2010). qI would affect the rate of oxygen evolution and therefore CO_2_ assimilation (Nawrocki et al. 2022). We observed however in the absence of PsbS or VDE, hence qE, in both aspen and *Arabidopsis* significant sustained quenching and NESD but only a neglectable effect on CO_2_ assimilation (Fig. 4). This is inconsistent with the idea that the slowly arising component in *npq1* upon extended NPQ induction is strictly connected to qI (Nilkens et al., 2010). Despite early studies reported a strong photoinhibition in *npq4* and *npq1* (Havaux and Niyogi, 1999; Li et al., 2002), under our experimental conditions with only red-actinic light for 80 minutes we did not observe an obvious decrease in CO_2_ assimilation during the rise of the sustained quenching, hence the increase in sustained quenching in unlikely to be solely due to increasing qI. In contrast, *cao2* showed a 75% decrease in CO_2_ assimilation as well as Y(II) decline, and large amount of sustained quenching without the production of NES supporting the suggestion of Nawrocki et al. (2022) that qI takes place at PSII RC without any spectral signatures. Moreover, by using *curt1*, impaired in thylakoid plasticity, we found that thylakoid reorganization is important for a proper response to increasing light (Garty et al., 2024; Nanda et al., 2025). In *curt1*, this impairment was associated with a marked decline in CO_2_ assimilation and Y(II), elevated levels of NESD, and a slow induction of sustained NPQ (Fig 4 H-K). Interestingly, stomatal conductance was affected in *curt1*, likely associated to an over-reduced PQ pool (Supplementary Fig. 14), a proposed cue for stomatal closure (Wang et al., 2016).

Although Y(NO) remains a poorly understood parameter, it simply reflects the relation between the minimal yield of fluorescence at any given time and its maximal in the dark-adapted state (Eq. 7). Thus, any increase in Y(NO) between the dark-relaxation phase and the dark-adapted state, reflects changes at the minimal yield of fluorescence product of impaired photochemistry, suggesting the presence of inactive broken PSII RC (Nawrocki et al., 2022). In HL treatments, we observed that after 20 minutes of dark-relaxation, Y(NO) was substantially increased in *cao2*, while it remained unchanged in *curt1*. Here, sustained increase in Y(NO) in *cao2* reflects the development of qI during HL exposure. In contrast photoinhibition may be partially or even entirely uncoupled from qI in *curt1*.

We observed a large amount of sustained quenching in classical *npq* mutants after prolonged exposure to HL evoking the original conclusions about *npq4* from Johnson and Ruban (2011). Thus, PsbS and Zea are most likely essential to rapidly fine-tune the rate of energy transfer from the antenna to the RC upon NPQ, but its requirement is not mandatory. In our work, we found that Y(II) decline represents a proxy of photoinhibition development during long NPQ inductions (Fig. 4). Previous studies have reported that Y(II) remains unaffected under increasing light intensities in both *npq1* and *npq4* mutants, even under HL treatments like those applied in this study (Külheim and Jansson, 2005; Ware et al., 2015). Conversely, no changes in CO_2_ assimilation were observed in Col-0 and *npq4* plants subjected to cycles of 100/1000 μE light fluctuations and no effects was found on biomass production under several fluctuating light conditions (Schiphorst et al., 2023).

Under natural conditions, *npq1* and *npq4* mutants can cope with light fluctuations with moderate effects at fitness level (Külheim et al., 2002) whereas the absence of other regulatory processes involved in photosynthetic control may have more drastic effects, Proton Gradient Regulator 5 involved in regulation of cyclic electron flow is for example more important (Soursa et al., 2012). Mutants lacking *curt1* possesses the fast photoprotective mechanisms associated with LHCII, but the absence of thylakoid plasticity results in reduced long-term responses to HL and fluctuating light environments (Pribil et al., 2018). This suggests that an interaction between NPQ and thylakoid rearrangements is necessary for plants to cope with changing light intensities. What remains to be understood is the relevance of thylakoid structural re-arrangements and to what extent the lateral heterogeneity applies in the light-adapted state where photosynthesis takes place (Andersson & Anderson, 1980; Garty et al., 2024; Nanda et al., 2025).

Taken together, our results suggests that slow photoprotective subprocesses in *npq* mutants are also dependent of LHCII and thylakoid re-arrangements, and PsbS and Zea are catalysts of these subprocesses instead of direct quenchers (Fig. 5 B). Moreover, the presence of NESD under photoinhibitory conditions in the absence of PsbS and Zea, makes this phenomenon dependent of LHCB proteins, which can be rapidly triggered by the above-mentioned regulators (Fig. 5 B). qH has been suggested to be a form of sustained photoprotective mechanism dependent on LHCII upon long actinic light exposure (Bru *et al*., 2022). Maybe qE, qZ, and qH and other potential photoprotective mechanisms – including spillover from PSII to PSI (Bag et al. 2020) – should be viewed not as separate mechanisms with distinct targets, but as overlapping components of a – yet not understood – holistic regulatory process that collectively modulate the functional antenna size of PSII, primarily by acting at the LHCB complexes. Finally, does slowly relaxing – sustained – quenching involving antenna component allow plants to cope with HL and/or does it in light-fluctuating conditions produce a fitness constraint that could been targeted to enhance photosynthesis in dynamic environments (Kromdijk et al., 2016; De Souza et al., 2022)? Comparative analyses of Arabidopsis and aspen mutants in natural conditions in combination with spectrally-time resolved fluorescence and gas exchange analysis may be the way to get additional insights about photosynthetic dynamics.

## Acknowledgements

We would like to thank Prof. Matthew P. Johnson and Alfred R. Holzwarth for their valuable inputs. We are grateful to Prof. Dario Leister and Dr. Anja Scheider (Ludwig-Maximilians-University Munich) for providing us with *curt1* seeds. The authors thank the work from the Poplar Transgenics Facility and the Wallenberg Lab Greenhouse. The presented research was supported by funding from the Swedish Foundation for Strategic Research (FFF20-0008) and the Swedish Research Council VR, Kempestiftelserna and Knut and Alice Wallenberg Foundation.

## References

Armbruster, U., Labs, M., Pribil, M., Viola, S., Xu, W., Scharfenberg, M., Hertle, A. P., Rojahn, U., Jensen, P. E., & Rappaport, F. (2013). Arabidopsis CURVATURE THYLAKOID1 proteins modify thylakoid architecture by inducing membrane curvature. The Plant Cell, 25(7), 2661–2678.

André, D., Marcon, A., Lee, K. C., Goretti, D., Zhang, B., Delhomme, N., Schmid, M., & Nilsson, O. (2022). FLOWERING LOCUS T paralogs control the annual growth cycle in Populus trees. Current Biology, 32(13), 2988-2996. e2984.

Andersson, B., & Anderson, J. M. (1980). Lateral heterogeneity in the distribution of chlorophyll-protein complexes of the thylakoid membranes of spinach chloroplasts. Biochimica et Biophysica Acta (BBA)-Bioenergetics, 593(2), 427–440.

Bag, P., Chukhutsina, V., Zhang, Z., Paul, S., Ivanov, A. G., Shutova, T., Croce, R., Holzwarth, A. R., & Jansson, S. (2020). Direct energy transfer from photosystem II to photosystem I confers winter sustainability in Scots Pine. Nature Communications, 11(1), 6388.

Bag, P. (2021). Light harvesting in fluctuating environments: evolution and function of antenna proteins across photosynthetic lineage. Plants, 10(6), 1184.

Baker, N. R. (2008). Chlorophyll fluorescence: a probe of photosynthesis in vivo. Annu. Rev. Plant Biol., 59(1), 89–113.

Bru, P., Steen, C. J., Park, S., Amstutz, C. L., Sylak-Glassman, E. J., Lam, L., Fekete, A., Mueller, M. J., Longoni, F., & Fleming, G. R. (2022). The major trimeric antenna complexes serve as a site for qH-energy dissipation in plants. Journal of Biological Chemistry, 298(11).

Caffarri, S., Kouřil, R., Kereïche, S., Boekema, E. J., & Croce, R. (2009). Functional architecture of higher plant photosystem II supercomplexes. The EMBO journal, 28(19), 3052–3063.

Carriquiry, M. A., Du, X., & Timilsina, G. R. (2011). Second generation biofuels: Economics and policies. Energy policy, 39(7), 4222–4234.

Chukhutsina, V. U., Holzwarth, A. R., & Croce, R. (2019). Time-resolved fluorescence measurements on leaves: principles and recent developments. Photosynthesis Research, 140(3), 355–369.

Croce, R., Canino, G., Ros, F., & Bassi, R. (2002). Chromophore organization in the higher-plant photosystem II antenna protein CP26. Biochemistry, 41(23), 7334–7343.

Croce, R., & van Amerongen, H. (2020). Light harvesting in oxygenic photosynthesis: Structural biology meets spectroscopy. Science, 369(6506), eaay2058.

Cutolo, E. A., Guardini, Z., Dall’Osto, L., & Bassi, R. (2023). A paler shade of green: engineering cellular chlorophyll content to enhance photosynthesis in crowded environments. New Phytologist, 239(5), 1567–1583.

De Souza, A. P., Burgess, S. J., Doran, L., Hansen, J., Manukyan, L., Maryn, N., Gotarkar, D., Leonelli, L., Niyogi, K. K., & Long, S. P. (2022). Soybean photosynthesis and crop yield are improved by accelerating recovery from photoprotection. Science, 377(6608), 851–854.

Engineer, C. B., Hashimoto-Sugimoto, M., Negi, J., Israelsson-Nordström, M., Azoulay-Shemer, T., Rappel, W. J., … & Schroeder, J. I. (2016). CO2 sensing and CO2 regulation of stomatal conductance: advances and open questions. Trends in plant science, 21(1), 16–30.

Farooq, S., Chmeliov, J., Wientjes, E., Koehorst, R., Bader, A., Valkunas, L., Trinkunas, G., & van Amerongen, H. (2018). Dynamic feedback of the photosystem II reaction centre on photoprotection in plants. Nature Plants, 4(4), 225–231.

Garab, G., Magyar, M., Sipka, G., & Lambrev, P. H. (2023). New foundations for the physical mechanism of variable chlorophyll a fluorescence. Quantum efficiency versus the light-adapted state of photosystem II. Journal of Experimental Botany, 74(18), 5458–5471.

Garty, Y., Bussi, Y., Levin-Zaidman, S., Shimoni, E., Kirchhoff, H., Charuvi, D., Nevo, R., & Reich, Z. (2024). Thylakoid membrane stacking controls electron transport mode during the dark-to-light transition by adjusting the distances between PSI and PSII. Nature Plants, 10(3), 512–524.

Genty, B., Briantais, J.-M., & Baker, N. R. (1989). The relationship between the quantum yield of photosynthetic electron transport and quenching of chlorophyll fluorescence. Biochimica et Biophysica Acta (BBA)-General Subjects, 990(1), 87–92.

Havaux, M., & Tardy, F. (1997). Thermostability and photostability of photosystem II in leaves of the Chlorina-f2 barley mutant deficient in light-harvesting chlorophyll a/b protein complexes. Plant physiology, 113(3), 913–923.

Havaux, M., & Niyogi, K. K. (1999). The violaxanthin cycle protects plants from photooxidative damage by more than one mechanism. Proceedings of the National Academy of Sciences, 96(15), 8762–8767.

Holzwarth, A. R., Miloslavina, Y., Nilkens, M., & Jahns, P. (2009). Identification of two quenching sites active in the regulation of photosynthetic light-harvesting studied by time-resolved fluorescence. Chemical Physics Letters, 483(4-6), 262–267.

Horton, P., Ruban, A., Rees, D., Pascal, A., Noctor, G., & Young, A. (1991). Control of the lightharvesting function of chloroplast membranes by aggregation of the LHCII chlorophyll—protein complex. FEBS letters, 292(1-2), 1–4.

Horton, P., Wentworth, M., & Ruban, A. (2005). Control of the light harvesting function of chloroplast membranes: the LHCII-aggregation model for non-photochemical quenching. FEBS letters, 579(20), 4201–4206.

Järvi, S., Suorsa, M., Paakkarinen, V., & Aro, E.-M. (2011). Optimized native gel systems for separation of thylakoid protein complexes: novel super-and mega-complexes. Biochemical Journal, 439(2), 207–214.

Johnson, M. P., & Ruban, A. V. (2009). Photoprotective energy dissipation in higher plants involves alteration of the excited state energy of the emitting chlorophyll (s) in the light harvesting antenna II (LHCII). Journal of Biological Chemistry, 284(35), 23592–23601.

Johnson, M. P., & Ruban, A. V. (2011). Restoration of rapidly reversible photoprotective energy dissipation in the absence of PsbS protein by enhanced ΔpH. Journal of Biological Chemistry, 286(22), 19973–19981.

Klughammer, C., & Schreiber, U. (2008). Complementary PS II quantum yields calculated from simple fluorescence parameters measured by PAM fluorometry and the Saturation Pulse method. PAM application notes, 1(2), 201–247.

Kromdijk, J., Głowacka, K., Leonelli, L., Gabilly, S. T., Iwai, M., Niyogi, K. K., & Long, S. P. (2016). Improving photosynthesis and crop productivity by accelerating recovery from photoprotection. Science, 354(6314), 857–861.

Kulheim, C., Ågren, J., & Jansson, S. (2002). Rapid regulation of light harvesting and plant fitness in the field. Science, 297(5578), 91–93.

Külheim, C., & Jansson, S. (2005). What leads to reduced fitness in non photochemical quenching mutants? Physiologia Plantarum, 125(2), 202–211.

Lam, L., Patel-Tupper, D., Lam, H. E., Steen, C. J., Ma, A., Ma, S. A., Leipertz, A., Lee, T.-Y., Fleming, G., & Niyogi, K. K. (2024). Resolving an unconventional non-photochemical quenching signature at the light-to-dark transition. bioRxiv, 2024.2010.2017.618902.

Leister, D. (2023). Enhancing the light reactions of photosynthesis: Strategies, controversies, and perspectives. Molecular Plant, 16(1), 4–22.

Li, X.-P., Müller-Moulé, P., Gilmore, A. M., & Niyogi, K. K. (2002). PsbS-dependent enhancement of feedback de-excitation protects photosystem II from photoinhibition. Proceedings of the National Academy of Sciences, 99(23), 15222–15227.

Malnoë, A. (2018). Photoinhibition or photoprotection of photosynthesis? Update on the (newly termed) sustained quenching component qH. Environmental and Experimental Botany, 154, 123–133.

Malnoë, A., Schultink, A., Shahrasbi, S., Rumeau, D., Havaux, M., & Niyogi, K. K. (2018). The plastid lipocalin LCNP is required for sustained photoprotective energy dissipation in Arabidopsis. The Plant Cell, 30(1), 196–208.

Nanda, S., Cainzos, M., Shutova, T., Fataftah, N., Fleig, V., Lihavainen-Bag, J., Bag, P., Holzwarth, A. R., & Jansson, S. (2025). Spillover is the dominant non-photochemical quenching mechanism in angiosperms. bioRxiv, 2025.2001.2026.634902.

Nanda, S., Shutova, T., Cainzos, M., Hu, C., Sasbrink, B., Bag, P., Blanken, T. d., Buijs, R., Gracht, L. v. d., & Hendriks, F. (2024). ChloroSpec: A new in vivo chlorophyll fluorescence spectrometer for simultaneous wavelength and time resolved detection. Physiologia Plantarum, 176(2), e14306.

Nawrocki, W. J., Liu, X., Raber, B., Hu, C., De Vitry, C., Bennett, D. I., & Croce, R. (2021). Molecular origins of induction and loss of photoinhibition-related energy dissipation qI. Science Advances, 7(52), eabj0055.

Nilkens, M., Kress, E., Lambrev, P., Miloslavina, Y., Müller, M., Holzwarth, A. R., & Jahns, P. (2010). Identification of a slowly inducible zeaxanthin-dependent component of non-photochemical quenching of chlorophyll fluorescence generated under steady-state conditions in Arabidopsis. Biochimica et Biophysica Acta (BBA)-Bioenergetics, 1797(4), 466–475.

Oxborough, K., & Baker, N. R. (1997). Resolving chlorophyll a fluorescence images of photosynthetic efficiency into photochemical and non-photochemical components–calculation of qP and Fv-/Fm-; without measuring Fo. Photosynthesis research, 54(2), 135–142.

Pribil, M., Sandoval-Ibáñez, O., Xu, W., Sharma, A., Labs, M., Liu, Q., Galgenmüller, C., Schneider, T., Wessels, M., & Matsubara, S. (2018). Fine-tuning of photosynthesis requires CURVATURE THYLAKOID1-mediated thylakoid plasticity. Plant physiology, 176(3), 2351–2364.

Ramakers, L. A., Harbinson, J., Wientjes, E., & van Amerongen, H. (2025). Unravelling the different components of nonphotochemical quenching using a novel analytical pipeline. New Phytologist, 245(2), 625–636.

Ruban, A. V. (2016). Nonphotochemical chlorophyll fluorescence quenching: mechanism and effectiveness in protecting plants from photodamage. Plant physiology, 170(4), 1903–1916.

Ruban, A. V., & Saccon, F. (2022). Chlorophyll a de-excitation pathways in the LHCII antenna. The Journal of Chemical Physics, 156(7).

Schiphorst, C., Koeman, C., Caracciolo, L., Staring, K., Theeuwen, T. P., Driever, S. M., Harbinson, J., & Wientjes, E. (2023). The effects of different daily irradiance profiles on Arabidopsis growth, with special attention to the role of PsbS. Frontiers in Plant Science, 14, 1070218.

Schreiber, U., Schliwa, U., & Bilger, W. (1986). Continuous recording of photochemical and non-photochemical chlorophyll fluorescence quenching with a new type of modulation fluorometer. Photosynthesis Research, 10, 51–62.

Suorsa, M., Järvi, S., Grieco, M., Nurmi, M., Pietrzykowska, M., Rantala, M., Kangasjärvi, S., Paakkarinen, V., Tikkanen, M., & Jansson, S. (2012). PROTON GRADIENT REGULATION5 is essential for proper acclimation of Arabidopsis photosystem I to naturally and artificially fluctuating light conditions. The Plant Cell, 24(7), 2934–2948.

Van Der Maaten, L., Postma, E., & Van den Herik, J. (2009). Dimensionality reduction: a comparative. J Mach Learn Res, 10(66-71).

Wang, W. H., He, E. M., Chen, J., Guo, Y., Chen, J., Liu, X., & Zheng, H. L. (2016). The reduced state of the plastoquinone pool is required for chloroplast mediated stomatal closure in response to calcium stimulation. The Plant Journal, 86(2), 132–144.

Ware, M. A., Belgio, E., & Ruban, A. V. (2015). Comparison of the protective effectiveness of NPQ in Arabidopsis plants deficient in PsbS protein and zeaxanthin. Journal of Experimental Botany, 66(5), 1259–1270.

Wilson, S., & Ruban, A. V. (2020). Rethinking the influence of chloroplast movements on non-photochemical quenching and photoprotection. Plant physiology, 183(3), 1213–1223.

