## Supplementary File for "Sustained quenching not always means photoinhibition"

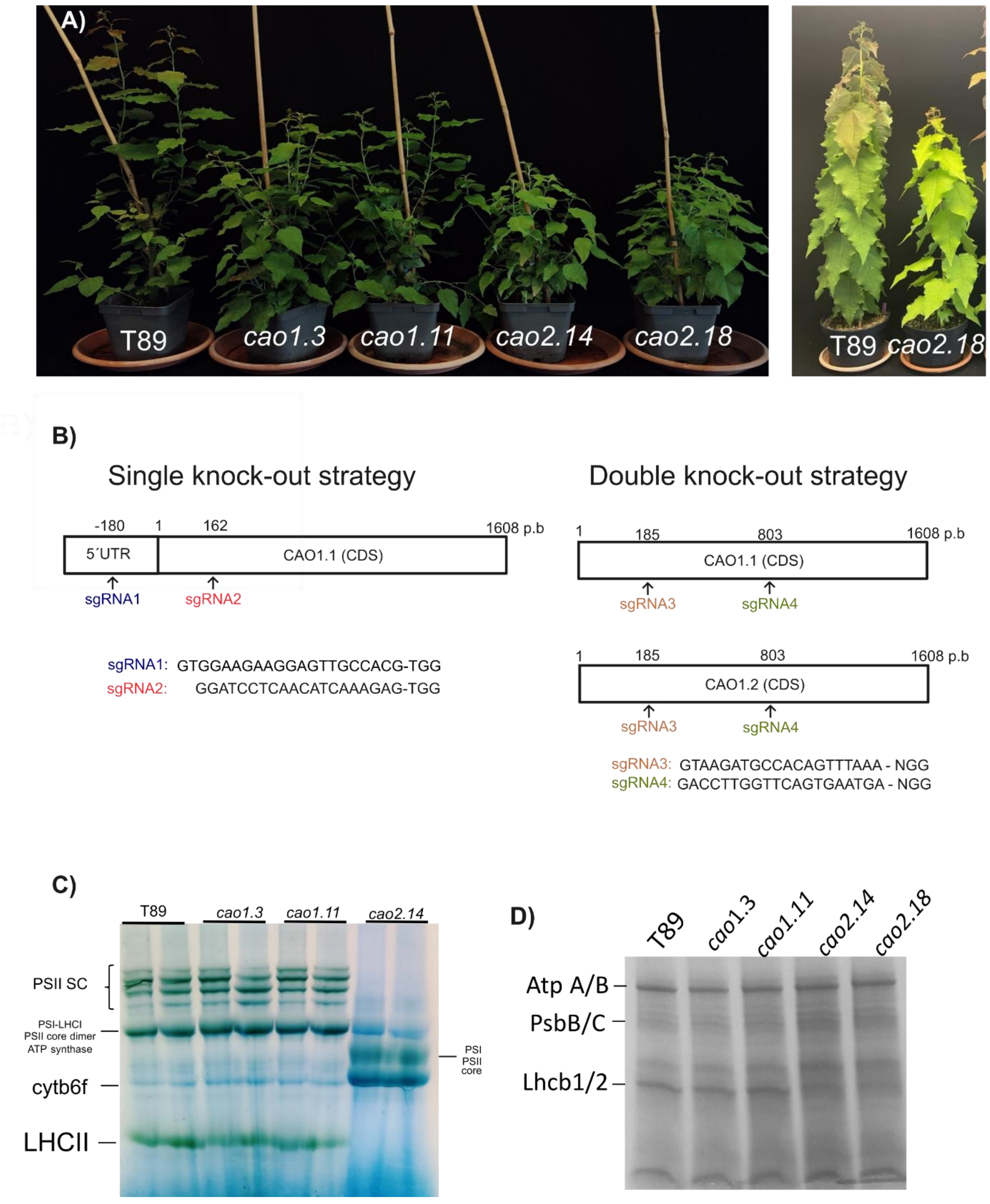


**Supplementary Figure 1. Collection of aspen chlorina mutants.** A) In the left panel, examples of young T89, *cao1.3*, *cao1.11*, *cao2.14* and *cao2.18* are shown. In the right panel, examples of adult T89 and *cao2.14* are shown. C) Molecular cloning design**.** sgRNA's for CAO1.1 and CAO1.1 + CAO1.2 were design to knock out one (*cao1*) or both genes (*cao2*). D) BN-PAGE from aspens thylakoids solubilized in 2% b-DM. E) SDS-PAGE from aspens thylakoids. 2 µg of chlorophylls were loaded per lane for each gel.

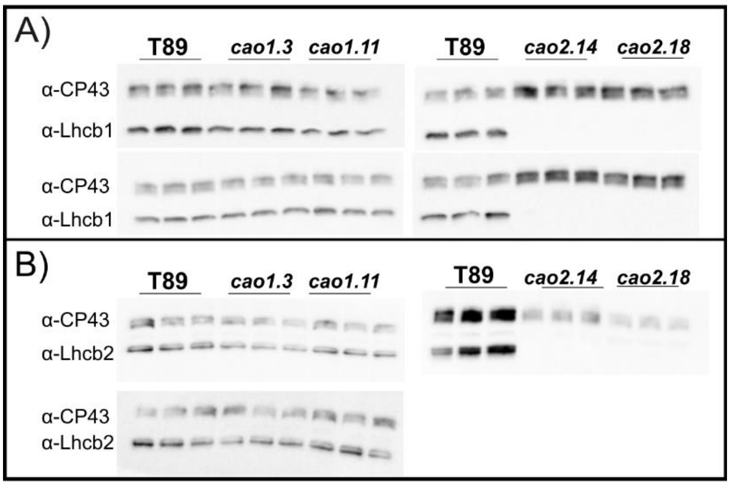


**Supplementary Figure 2. Immunoblots of Lhcb proteins in T89 and *cao* mutants.** A) 2 µg of chlorophylls were loaded into SDS-Page and blotted against α-CP43 and α-Lhcb1 for T89 reference line and chlorina mutants. B) 2 µg of were loaded into SDS-Page and blotted against α-CP43 and α-Lhcb2 for T89 reference line and chlorina mutants.

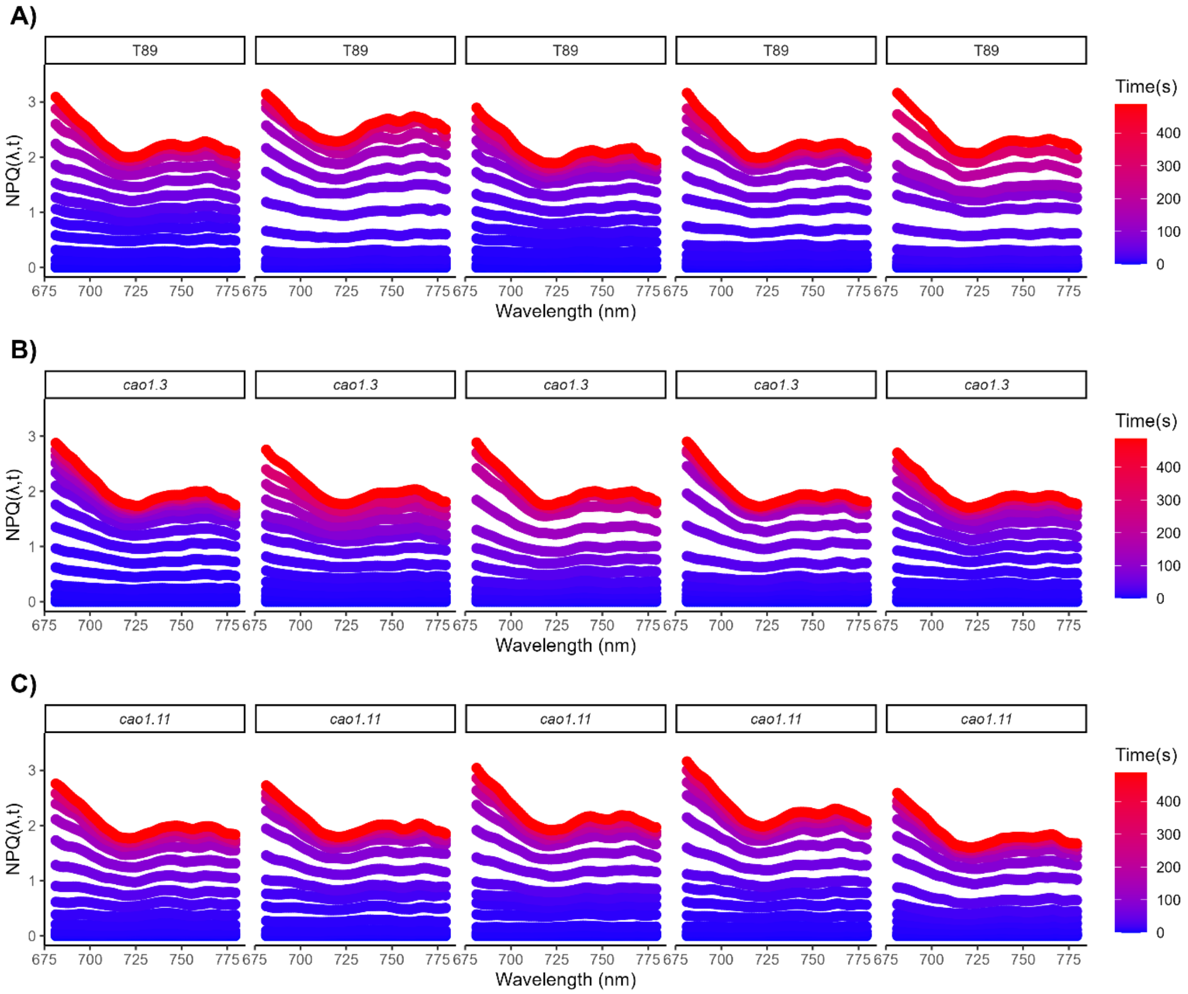


**Supplementary Figure 3. T89 and *cao1* NPQ evolution spectra.** T89 and *cao1* mutants were illuminated with 1000 µE. NPQ spectra was recorded as NPQ $(\lambda,t) = F(\lambda,t_0) / F(\lambda,t) - 1$. A), B), C) show NPQ evolution spectra from T89, *cao1.3* and *cao1.11*, respectively. Blue spectra are associated to the initial phase of the NPQ induction, whereas red spectra indicate quenched spectra after 8 minutes of actinic light exposure. Each NPQ evolution spectra represents one biological replicates. Data are from 5 biological replicates.

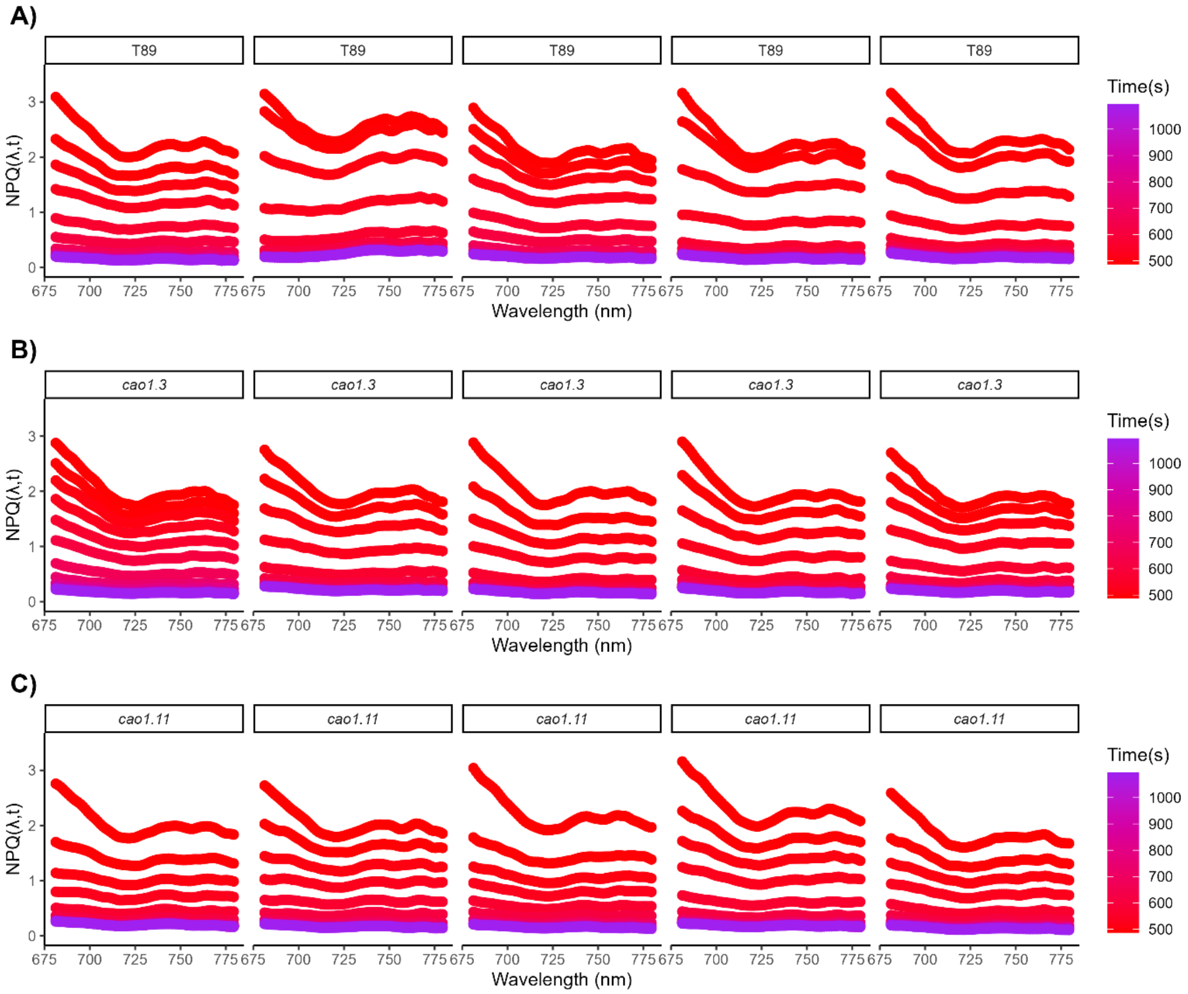


**Supplementary Figure 4. T89 and *cao1* relaxation spectra.** T89 and *cao1* mutants were illuminated with 1000 µE. NPQ spectra was recorded as NPQ (λ,t)= F (λ,$t_0$) / F (λ,t) - 1. A), B), C) show NPQ relaxation evolution spectra from T89, *cao1.3* and *cao1.11*, respectively. Red spectra are associated to the end of the initial phase of the NPQ induction, whereas purple spectra indicate spectra after 10 minutes of dark relaxation. Each NPQ evolution spectra represents one biological replicates. Data are from 5 biological replicates. Note that from the top quenched spectra measured at 487.5 s, the subsequent spectra were recorded at 493.1, 503.6 and 524.2 s in dark, where NESD is almost abolished.

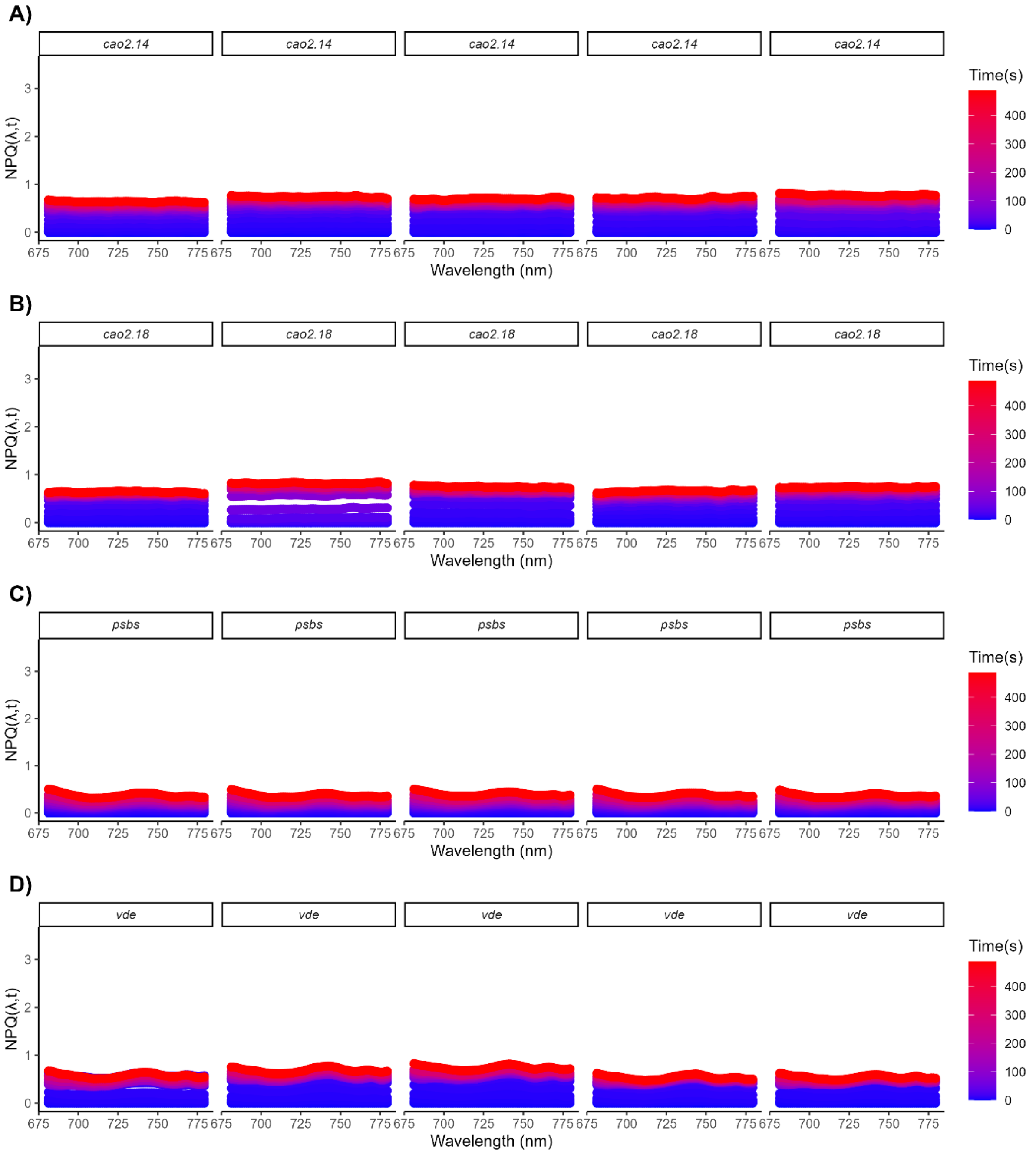


**Supplementary Figure 5. NPQ evolution spectra of NPQ mutants.** NPQ mutants, *cao2*, *psbs* and *vde* were illuminated with 1000 μE. NPQ spectra was recorded as NPQ (λ,t)= F (λ,$t_0$) / F (λ,t) - 1. A), B), C) and D), show NPQ evolution spectra from *cao2.14*, *cao2.18, psbs* and *vde*, respectively. Blue spectra are associated to the initial phase of the NPQ induction, whereas red spectra indicate quenched spectra after 8 minutes of actinic light exposure. Each NPQ evolution spectra represents one biological replicates. Data are from 5 biological replicates.

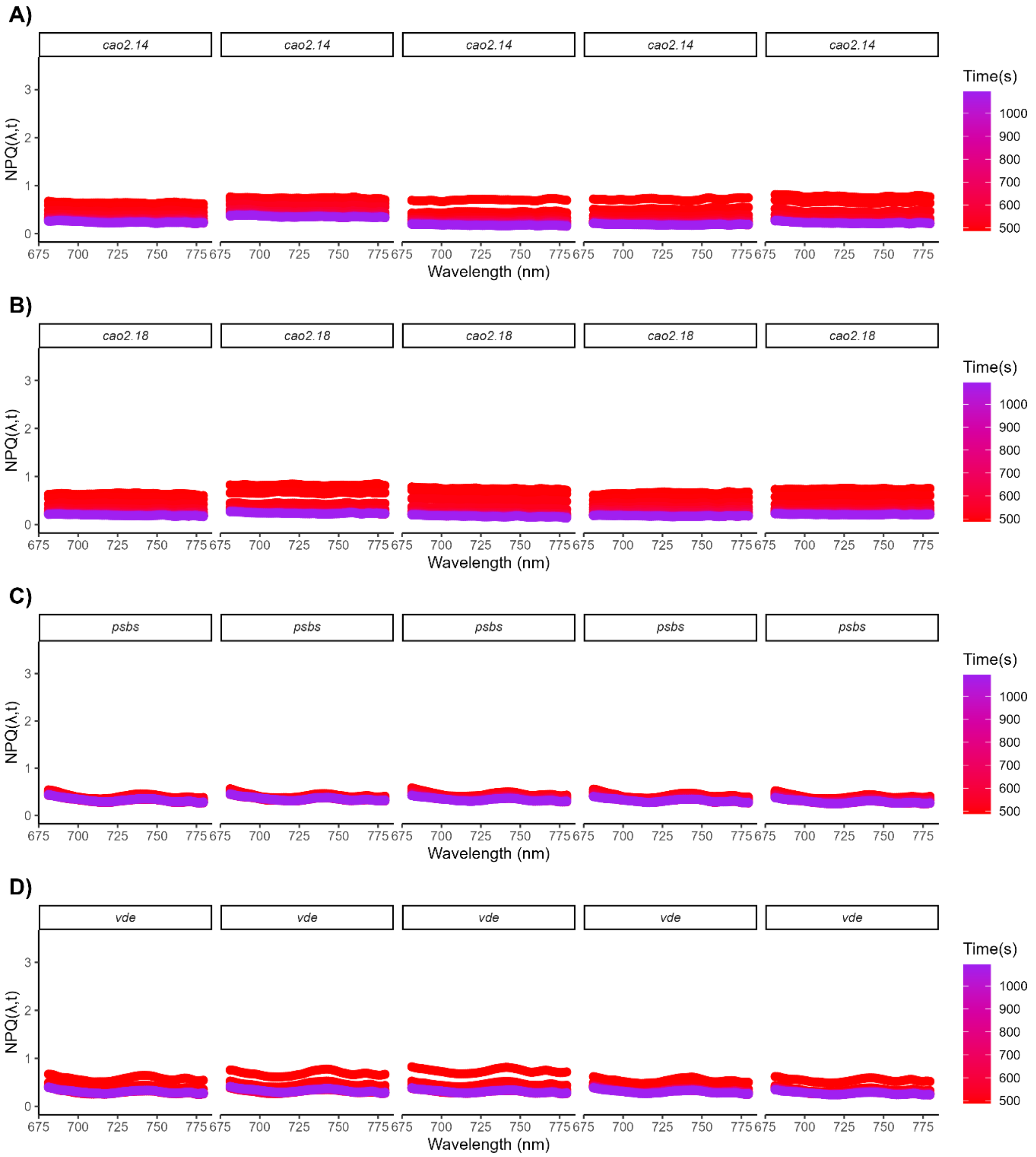


**Supplementary Figure 6. NPQ relaxation evolution spectra of NPQ mutants.** NPQ mutants, *cao2*, *psbs* and *vde* were illuminated with 1000 µE. NPQ spectra was recorded as NPQ $(\lambda,t) = F(\lambda,t_0) / F(\lambda,t) - 1$. A), B), C) and D), show NPQ relaxation evolution spectra from *cao2.14*, *cao2.18, psbs* and *vde*, respectively. Red spectra are associated to the end of the initial phase of the NPQ induction, whereas purple spectra indicate spectra after 10 minutes of dark relaxation. Each NPQ evolution spectra represents one biological replicates. Data are from 5 biological replicates.

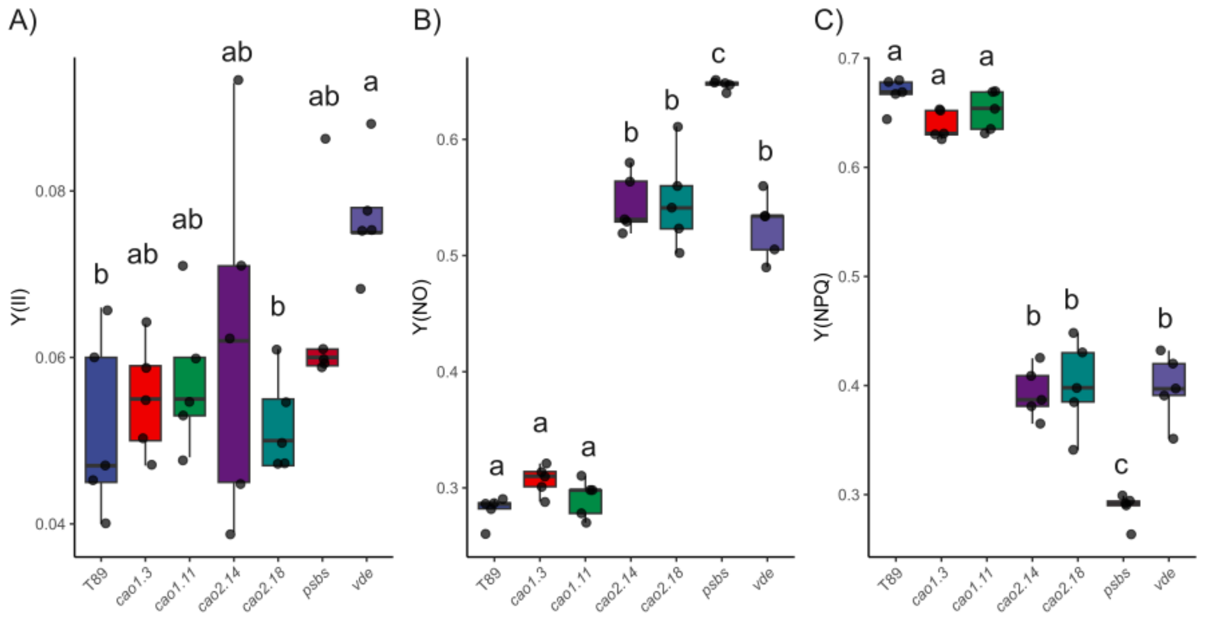


**Supplementary Figure 7. PSII energy partitioning from PAM fluorometry.** NPQ inductions were performed in aspens npq mutants and Y(II), Y(NO) and Y(NPQ) was determined in A), B and C), respectively. Data represents values achieved once T89 reached steady state at 460s of the NPQ induction. Each point indicates a different biological replica (n = 5). Shared letters between groups indicates non-significant differences according to Tukey’s test ($p < 0.05$).

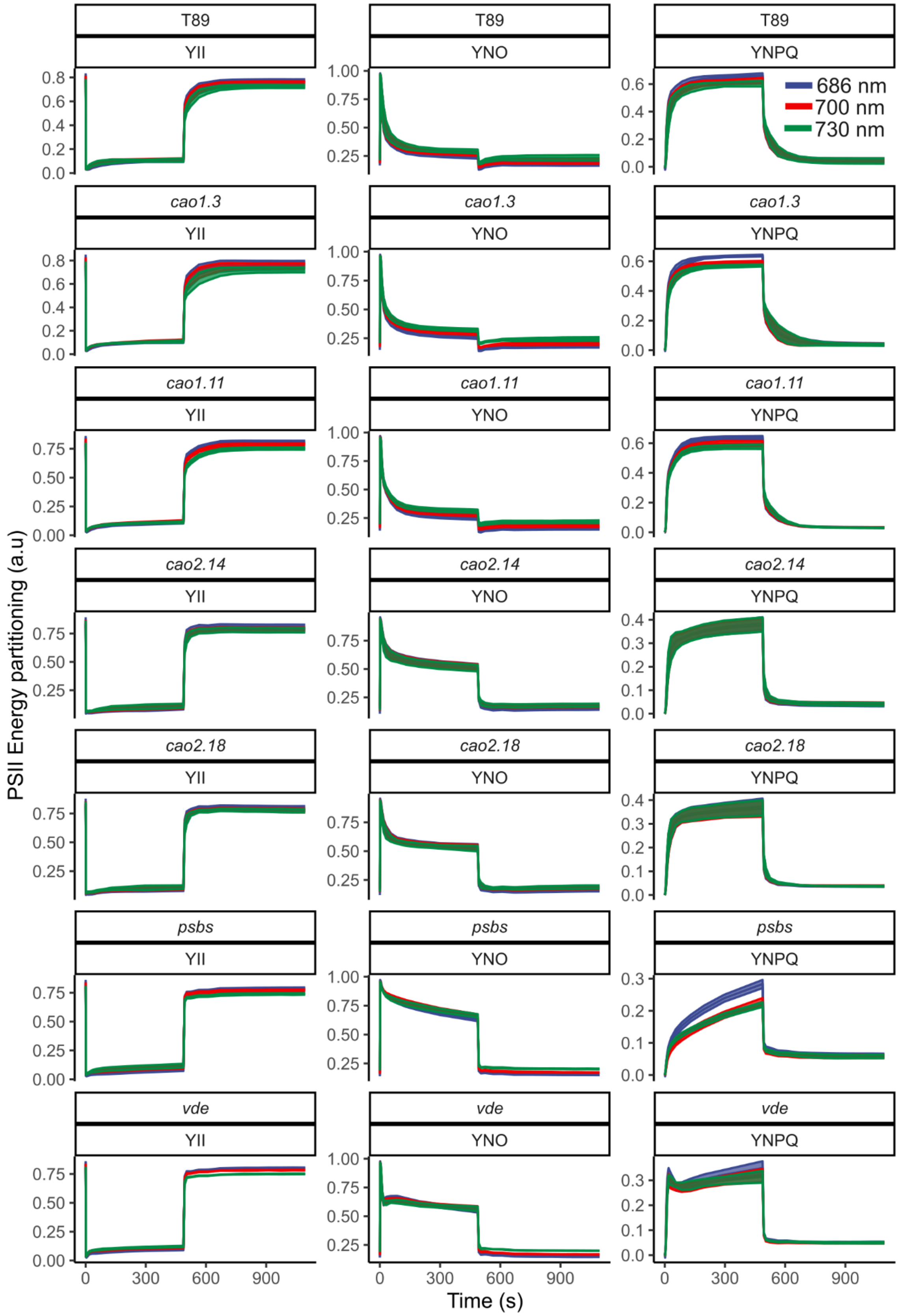


**Supplementary Figure 8. Apsens npq mutants PSII energy partitioning heterogeneity.** Y(II), Y(NPQ) and Y(NO) was monitored in T89 and *cao1, cao2, psbs and vde* upon NPQ inductions at 1000

µE. Each plot represents the values for genotype and the respective parameter indicated at the top of each trace. Data are mean ± s.d (n > 5 biologically independent experiments).

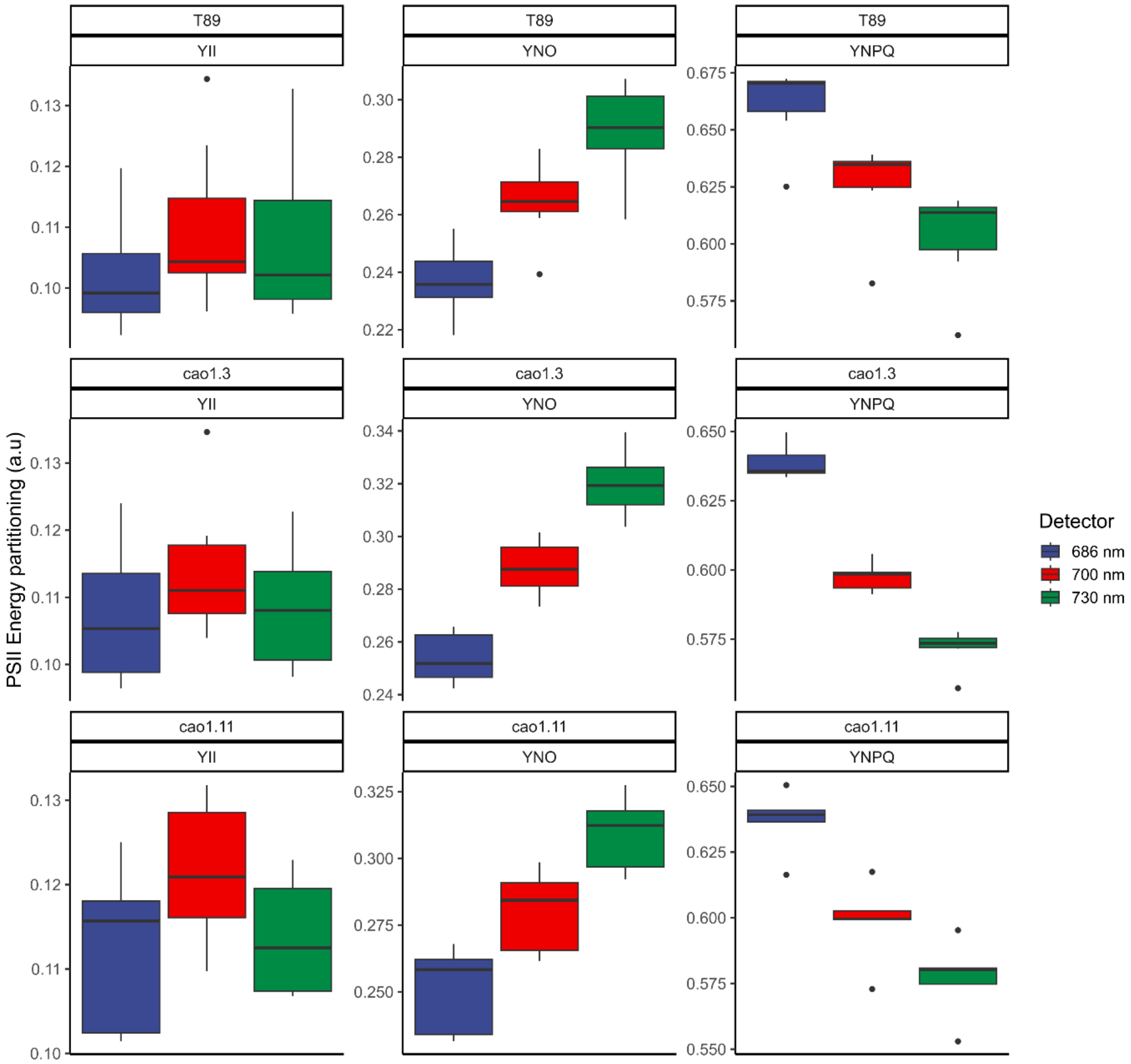


**Supplementary Figure 9. T89 and *cao1* PSII energy partitioning heterogeneity.** Y(II), Y(NPQ) and Y(NO) was monitored in T89 and *cao1* upon NPQ inductions at 1000 µE. At the top is indicated for each boxplot genotype and respective parameter for PSII energy partitioning measured at 487.5 s during the light induction phase. Data is from at least 5 independent biological replicas.

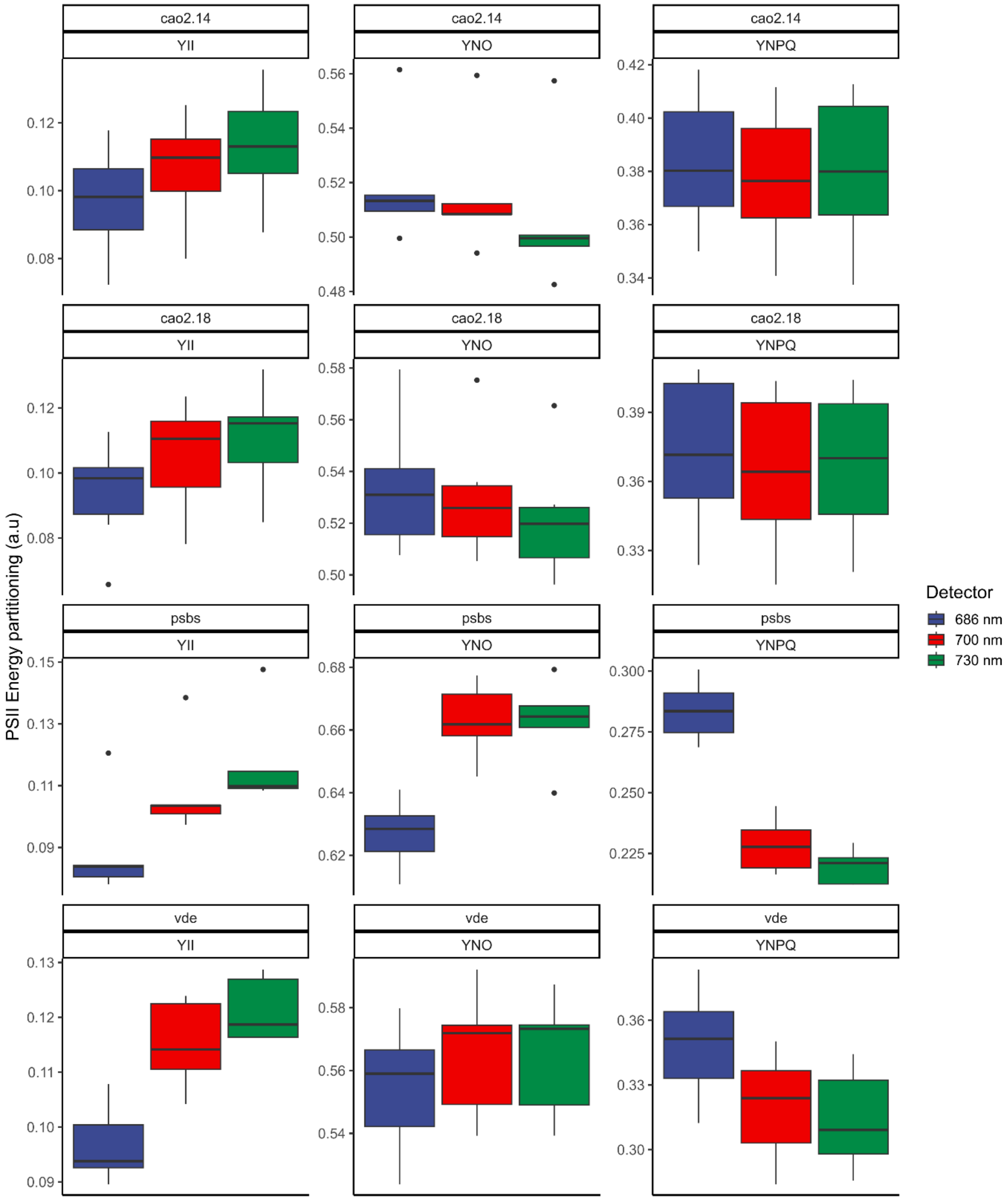


**Supplementary Figure 10. Aspen NPQ mutants PSII energy partitioning heterogeneity.** Y(II), Y(NPQ) and Y(NO) was monitored in *cao2*, *psbs* and *vde* upon NPQ inductions at 1000 µE. At the top is indicated for each boxplot genotype and respective parameter for PSII energy partitioning measured at 487.5 s during the light induction phase. Data is from at least 5 independent biological replicas.

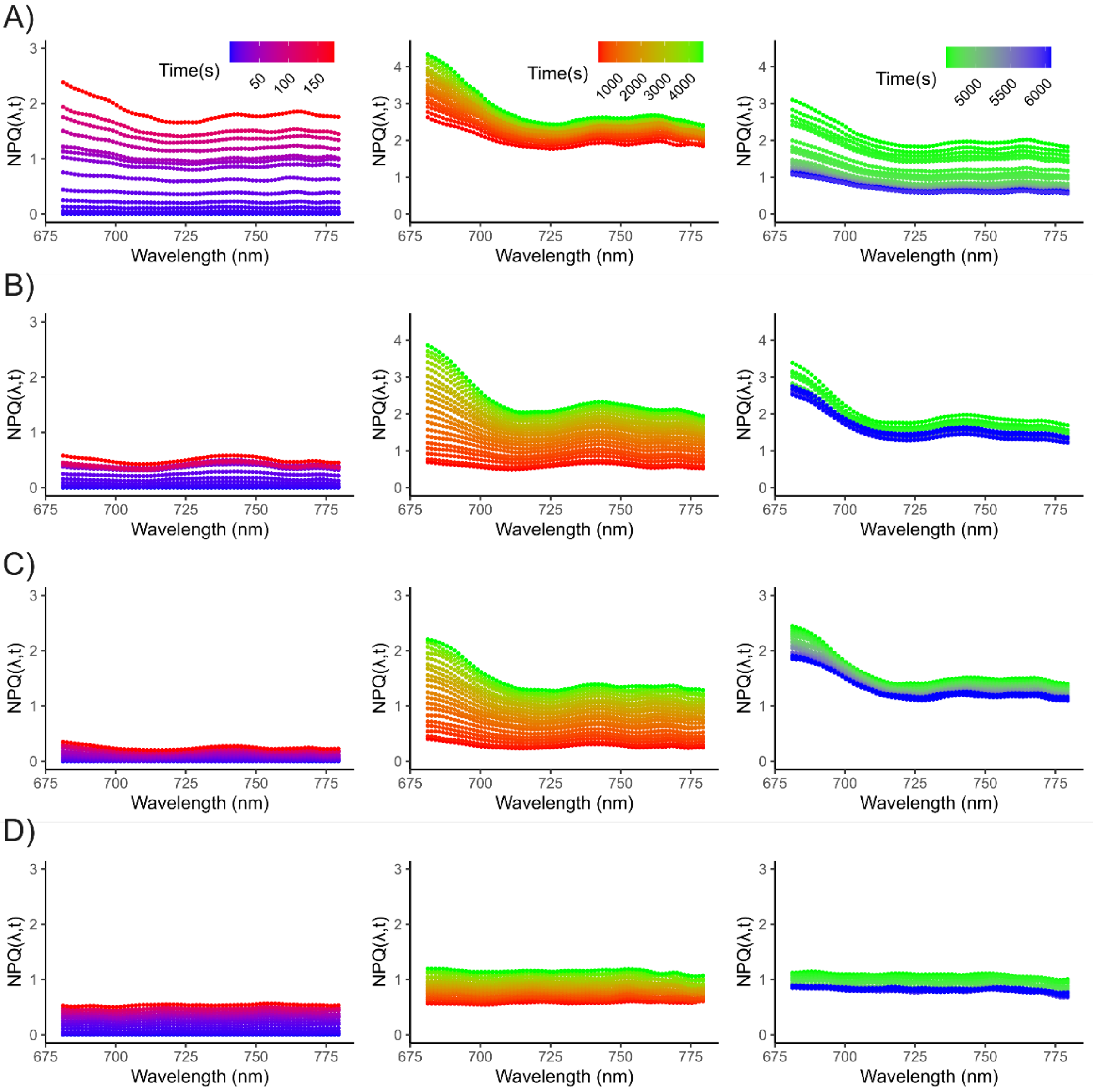


**Supplementary Figure 11. Sustained NPQ spectra of aspen npq mutants**. A), B), C), and D) show the NPQ spectra for T89, *vde*, *psbs*, and *cao2*, respectively. The left panels show the NPQ spectra during the first 200 seconds, highlighting the induction of qE and qZ. In T89, the spectra are highly heterogeneous, while the mutants show more homogeneous spectra. The middle panels show the development of NES over time, from 200s to 4600s. Notably, the spectra in *vde* and *psbs* become heterogeneous as sustained quenching develops. The right panels show the NPQ spectra during the relaxation phase. Sustained quenching is almost uniform in T89, but strongly heterogeneous in *vde* and *psbs.*

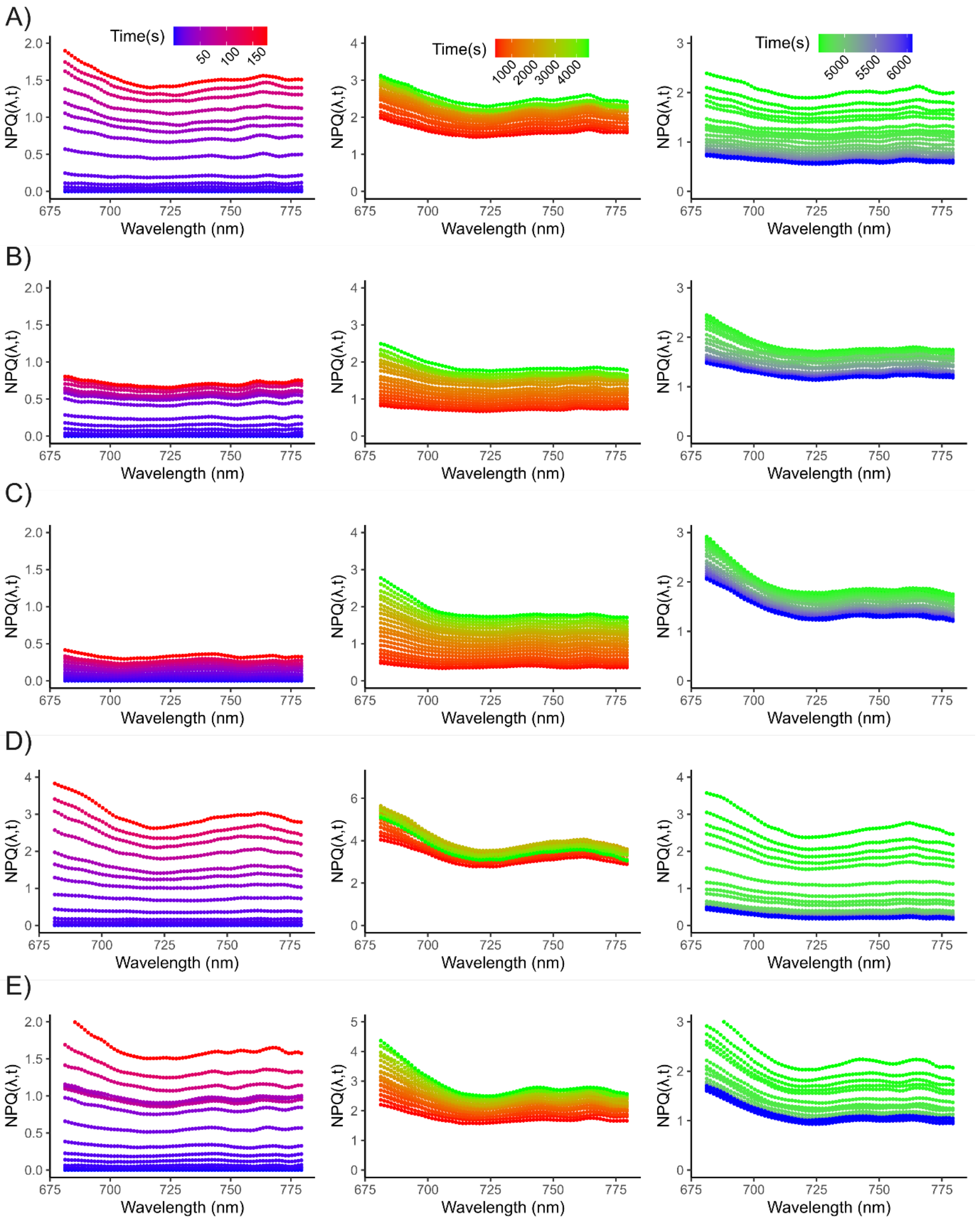


**Supplementary Figure 12. Sustained NPQ spectra of *Arabidopsis* npq mutants**. A), B), C), D) and E) show the NPQ spectra for Col-0, *npq1*, *npq4*, L17 and *curt1*, respectively. The left panels show the NPQ spectra during the first 200 seconds, highlighting the induction of qE and qZ. The middle panels show the development of NES over time, from 200s to 4600s. The right panels show the NPQ spectra during the relaxation phase.

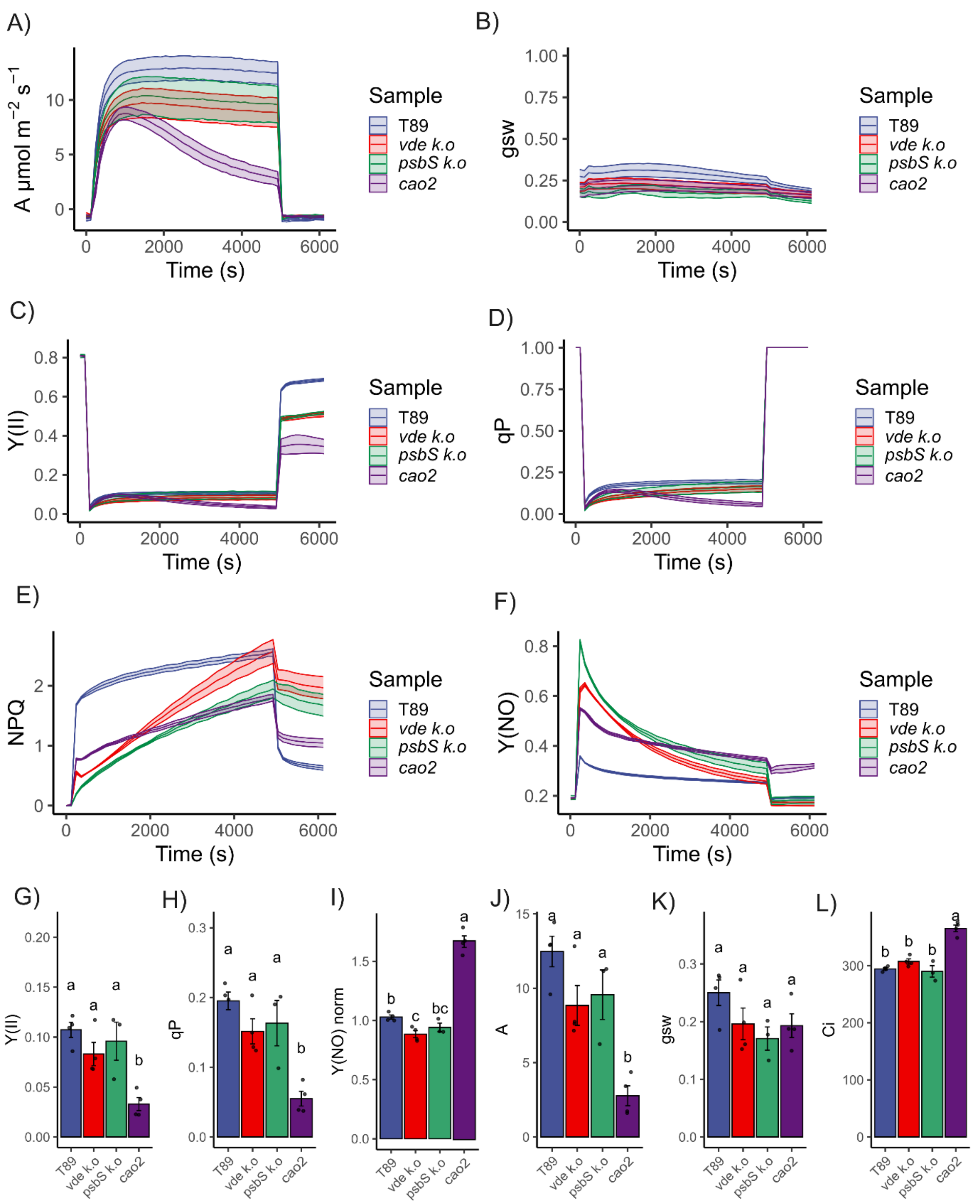


**Supplementary Figure 13. Photoinhibition and NPQ in aspen mutants after HL treatment.** NPQ kinetics were performed as described in Figure 5. A), B), C), D), E) and F) show A, stomatal conductance (gsw), Y(II), qP, NPQ and Y(NO) kinetics, respectively. Y(II) and A was normalized to their respective maximal between the range of 480s to 4920s and presented in Figure 5. Data are mean ± s.e (n > 3 biologically independent experiments). Note that *vde* at the end of the induction phase has even higher NPQ than T89 and in both *vde* and *psbs* a large amount of sustained quenching is developed which has little effect on $CO_2$ assimilation. Note that the signature of high Y(NO) decreases in time in *vde* and *psbs* mutants. Although *cao2* has a smaller absorption cross-section and lower chlorophyll content per area, assimilation declines up to 75% during HL treatments. At the relaxation phase, Y(NO) does not relax in *cao2* suggesting the presence of broken PSII RC. G), H), J), K) and L) show values at the end of the high light treatment for Y(II), qP, A, gsw and Ci, respectively. I) show values at the end of the dark-relaxation phase for Y(NO) normalized to Y(NO) at t = 0. Shared letters between groups indicates non-significant differences according to Tukey's test ($p < 0.05$). Each point represents a different biological replica.

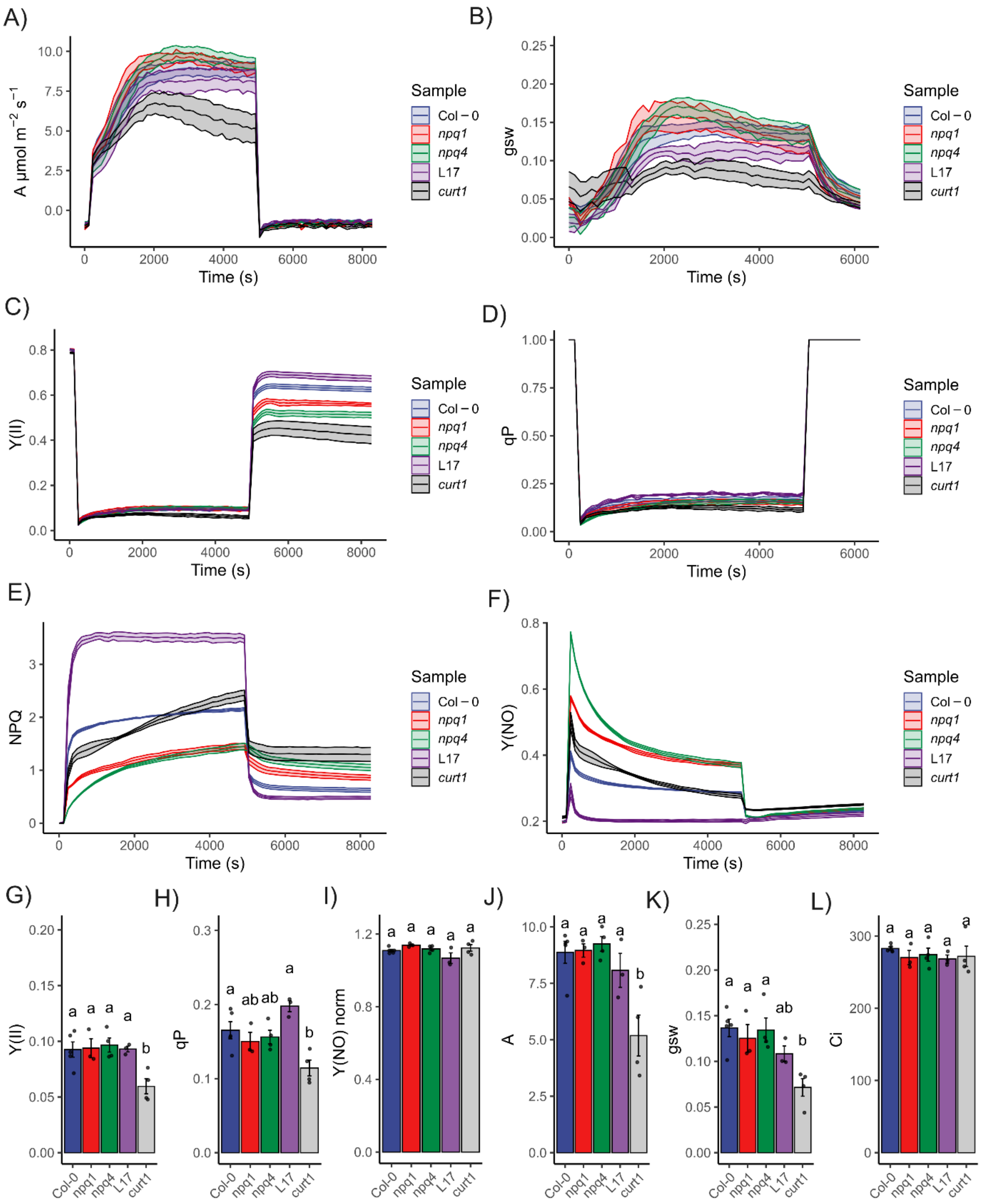


**Supplementary Figure 14. Photoinhibition is enhanced in the absence of thylakoid reorganization.** NPQ kinetics were performed at 1500 µE for 4680s with a relaxation phase of 3500s in *Arabidopsis* mutants. A), B), C), D), E) and F) show A, gsw, Y(II), qP, NPQ and Y(NO) kinetics, respectively. Y(II) and A was normalized to their respective maximal between the range of 480s to 4920s and presented in Figure 5. Data are mean ± s.e (n > 3 biologically independent experiments). Note that *npq1 and npq4* developed large amounts of sustained quenching. In *curt1* the decline in A was followed by Y(II). G), H), J), K) and L) show values at the end of the high light treatment for Y(II), qP, A, gsw and Ci, respectively. I) show values at the end of the dark relaxation interval for Y(NO) normalized to Y(NO) at t = 0. Shared letters between groups indicates non-significant differences according to Tukey's test ($p < 0.05$). Each point represents a different biological replica.

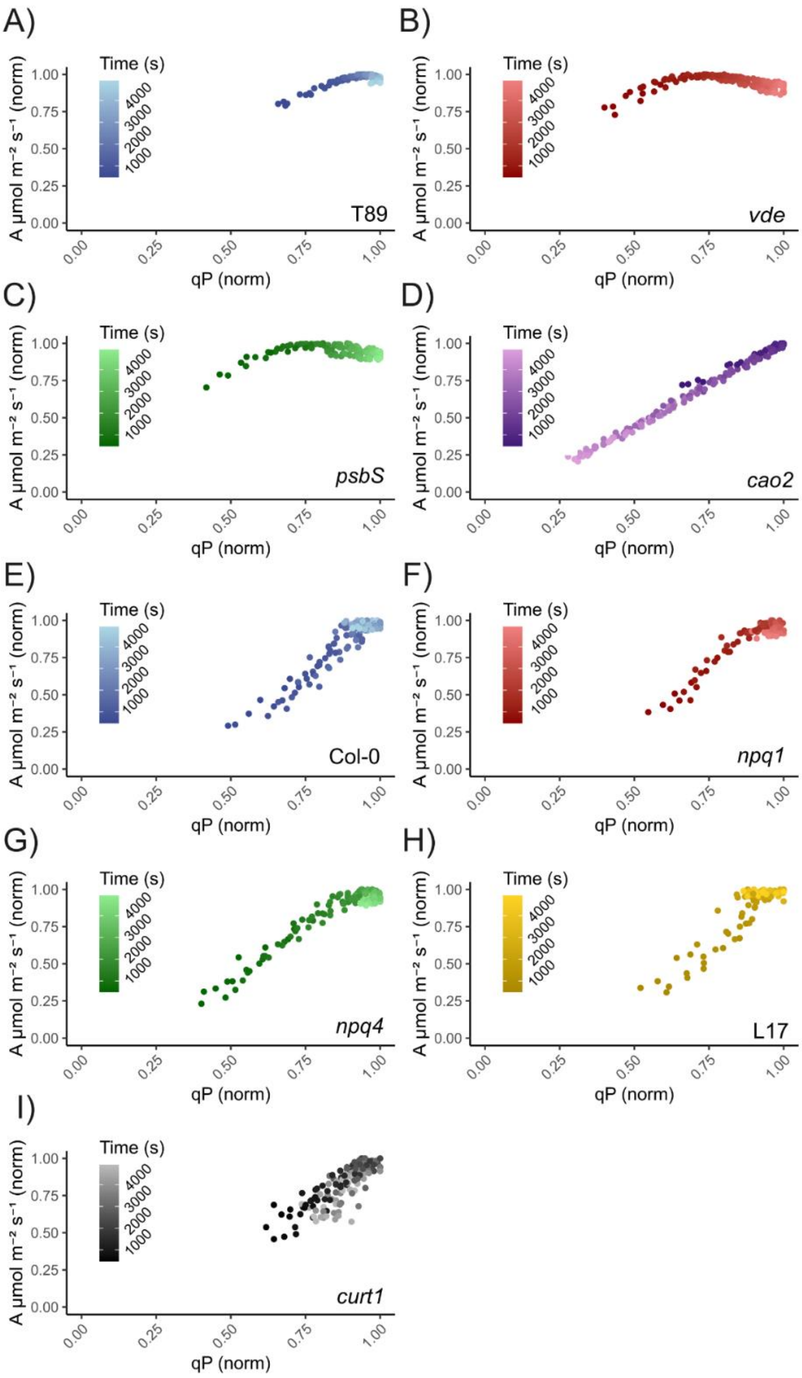


**Supplementary Figure 15. qP kinetics as a function of A.** Normalized A is shown as a function of normalized qP. A), B), C), D), E), F), G), H), I) show relations for T89, *vde, psbs, cao2,* Col-0, *npq1, npq4,* L17 and curt1, respectively. Note that very similar trend was found when Y(II) norm was compared to A (norm), although generally, lower normalized qP values were detected at earlier time points.

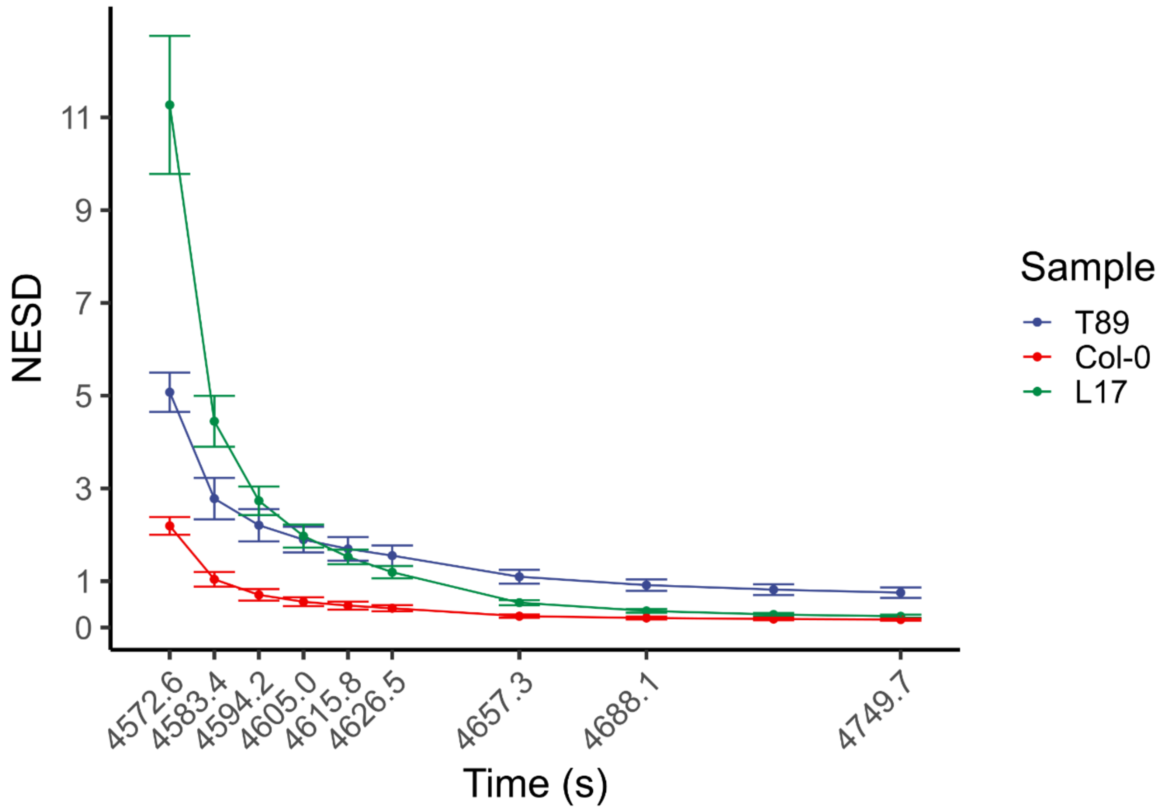


**Supplementary Figure 16. Sustained NES after photoinhibition.** NESD relaxation phase of NPQ kinetics for T89, Col-0 and L17 after HL treatment (1500 µE for 4572.6 s). Note that in Col-0 NES relax after 60 s whereas in L17 takes up to 110 s. In T89, NES are found after 180 s of dark relaxation.
